# Structural transitions in kinesin minus-end directed microtubule motility

**DOI:** 10.1101/2024.07.29.605428

**Authors:** Satoki Shibata, Matthew Y. Wang, Tsuyoshi Imasaki, Hideki Shigematsu, Yuanyuan Wei, Chacko Jobichen, Hajime Hagio, J. Sivaraman, Sharyn A. Endow, Ryo Nitta

**Author notes:** Equal contribution (Satoki Shibata, Matthew Y. Wang, Tsuyoshi Imasaki). Centre for Advanced Microscopy, The Australian National University, Canberra ACT 2601, AUS.

## Abstract

Kinesin motor proteins hydrolyze ATP to produce force for spindle assembly and vesicle transport, performing essential functions in cell division and motility, but the structural changes required for force generation are uncertain. We now report high-resolution structures showing new transitions in the kinesin mechanochemical cycle, including power stroke fluctuations upon ATP binding and a post-hydrolysis state with bound ADP + free phosphate. We find that rate-limiting ADP release occurs upon microtubule binding, accompanied by central β-sheet twisting, which triggers the power stroke – stalk rotation and neck mimic docking – upon ATP binding. Microtubule release occurs with β-strand-to-loop transitions, implying that β-strand refolding induces Pi release and the recovery stroke. The strained β-sheet during the power stroke and strand-to-loop transitions identify the β-sheet as the long-sought motor spring.

**Teaser:** Stalk rotation, β-sheet twisting and refolding, and neck mimic docking drive the reversed working stroke of kinesin-14

**INTRODUCTION:** Kinesin family proteins couple ATP hydrolysis to microtubule binding, generating force to produce steps or displacements along microtubules. The mechanism by which kinesins and other cytoskeletal motor proteins produce force is not fully understood. A current hypothesis is that the motors contain a spring-like or elastic element that creates strain under load during nucleotide binding or release, followed by a strain-relieving conformational change that produces force and a working stroke of the motor. The spring has not yet been identified for any motor. The power stroke differs for different motors – it consists of neck linker docking for plus-end directed kinesin-1 or a swing of the helical stalk for minus-end directed kinesin-14.

**RATIONALE:** Despite considerable research, the molecular dynamics of the kinesin-14 power stroke are still obscure, impeded by the weak microtubule binding of the motor. We overcame the weak binding by introducing a point mutation into the motor that results in faster ATP hydrolysis than wild type and tighter microtubule binding, which enabled us to resolve the motor mode of action. We now present high-resolution cryo-electron microscopy (cryo-EM) and x-ray structures of key mechanochemical states across the full force-producing cycle of a kinesin dimeric motor.

**RESULTS:** The new structures represent five different nucleotide states – two pre-power stroke states, a fluctuating power stroke, and two post-power stroke states. The structures are both microtubule-attached and unattached. They show the motor trapped in previously unreported transition states and reveal new conformational changes involved in energy transduction. The new transition states include a transient state in which the power stroke fluctuates during ATP binding and a new state of a kinesin motor bound to ADP and free Pi prior to phosphate release. The conformational changes include the folding of the kinesin-14 neck mimic into a structure resembling the docked kinesin-1 neck linker, accompanying the power stroke, and previously unreported β-strand-to-loop transitions with stored free energy that potentially induce Pi release and drive the recovery stroke. We interpret the new structures in the context of the hypothesis that the central β-sheet undergoes distortional changes during the mechanochemical cycle that store and release free energy, functioning as the elusive spring of the motors.

**CONCLUSION:** The new structures show that force is produced by coupled movements of the helical stalk, central β-sheet, and neck mimic, and uncover structural changes during the power stroke that are conserved among kinesins and myosin. We find that kinesin-14 binds to a microtubule by one head during the mechanical cycle, undergoes rate-limiting ADP release, and changes in conformation during ATP binding and hydrolysis to produce force. Notably, kinesin-14 utilizes the same mechanical strategy for force production as other kinesins but couples the changes to a large swing of the stalk, an innovation derived from myosin that is not observed for kinesin-1 or other kinesin motors. Force is produced by rearranging the binding surfaces of the stalk, strand β1, helices ɑ4 and ɑ6, and the neck mimic, and by twisting and shortening strands of the central β-sheet. These structural changes produce a power stroke – rotation of the helical stalk accompanied by neck mimic docking – during the transition from the nucleotide-free to ATP-bound state, and a reverse stroke after phosphate release that reprimes the motor for the next microtubule binding interaction.

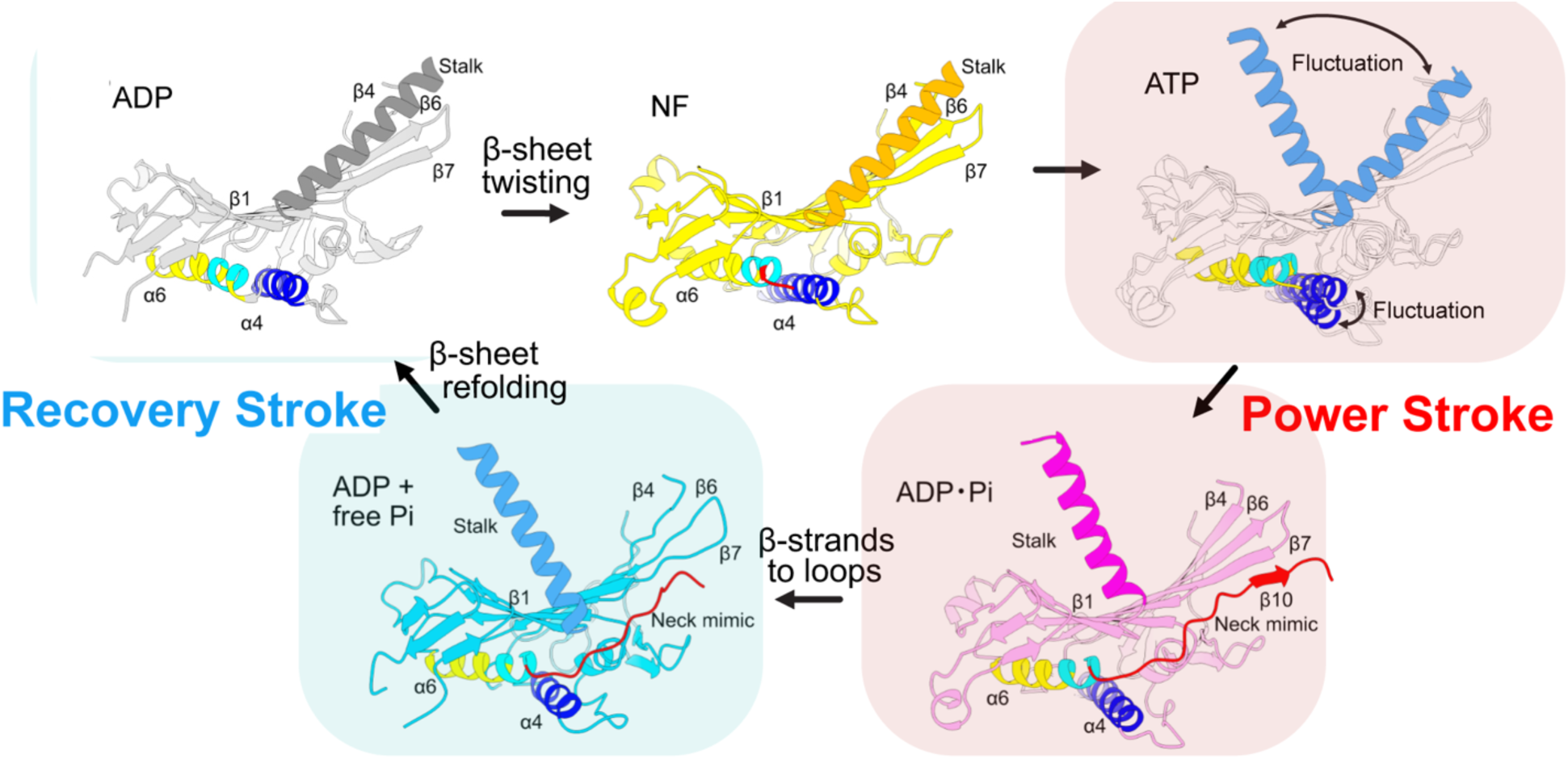

**Kinesin-14 force production:** New transition states and structural movements in a model for motor energy transduction and force production: β-sheet twisting stores free energy in the microtubule-bound nucleotide-free (NF) state. A fluctuating power stroke is produced in the ATP state with neck mimic docking in the ADP·Pi state, resembling the kinesin-1 neck linker. This is followed by β-strand-to-loop transitions in the microtubule-bound ADP + free Pi state. Finally, β-sheet refolding drives the recovery stroke for reversion to the ADP state.

## Main Text

Motor proteins – the dyneins, myosins, and kinesins – hydrolyze ATP and bind to microtubules or actin filaments, producing force and steps along their filament or sliding of filaments relative to one another. The mechanism by which motor proteins produce force and movement is still not fully understood. The motor force-generating mechanism is known to rely on ATP binding, hydrolysis, and release of hydrolysis products, coupled to filament binding and release (*1*). This has led to the idea that motility is regulated by the nucleotide state of the motor, together with motor-filament interactions (*2, 3*). Motors have been hypothesized to contain a spring-like or elastic element that compresses in a specific nucleotide state, storing free energy, and releases in a subsequent state, producing force (*1*), but the spring has not yet been identified. The movements in the motor domain due to nucleotide binding, hydrolysis, and product release are small, based on kinetic studies of myosin (*4*), a motor evolutionarily related to kinesins (*5*), and other ATP-binding enzymes (*6*). This has also been observed in structural studies of myosin in different nucleotide states (*7, 3*).

The kinesins produce a power stroke, but the structural element that functions as a lever to amplify force produced by the motor differs for kinesin-1 and kinesin-14, motors that move in opposite directions on microtubules. Kinesin-1, the first-discovered kinesin, is a highly processive motor that moves towards microtubule plus ends (*8–10*). The kinesin-1 power stroke is thought to consist of a large change in conformation of the neck linker, a short region (∼13 amino acids) that joins the conserved motor domain to the α-helical coiled-coil stalk (*11, 12*). Upon ATP binding, the neck linker binds to, or docks, onto the motor domain, but is mobile after ATP hydrolysis and Pi release. A second short region (∼11 amino acids) at the N terminus of the conserved motor domain, the cover strand, forms a β-strand (β0) that binds to the neck linker during force production (*11, 13*).

By contrast, kinesin-14 Ncd is a motor of low processivity that binds to microtubules and produces displacements towards the microtubule minus ends, differing from kinesin-1. The Ncd coiled-coil stalk in crystal structures of the dimeric motor shows a large ∼75-80° rotation towards the microtubule minus end (*14–17*), resembling the swing of the myosin lever arm (*18, 19*). A large displacement by Ncd observed in laser-trap assays and the dependence of motor velocity on the length of the stalk support the hypothesis that the Ncd stalk functions like a lever (*20, 14, 21*). The kinesin-14 stalk is thus thought to produce the working stroke of the motor. A short region (10-11 amino acids) at the kinesin-14 C terminus, the neck mimic, is also thought to contribute to the motor power stroke (*22, 23*). This region is analogous to the kinesin-1 neck linker (*24*). However, unlike the kinesin-1 neck linker, the kinesin-14 Ncd neck mimic is largely disordered in previous crystal structures.

Although basic features of force production by the kinesins have been reported, key steps in the motor mechanism are still not known, including the residues involved in mechanochemical coupling and the conformational changes that occur upon microtubule binding and nucleotide release, triggering the power stroke. Studies on myosin identified the α-helical light-chain binding region immediately adjacent to the conserved motor domain as a lever-like element that produces a power stroke by undergoing a large rotation, amplifying small movements of structural elements in the conserved motor domain (*19, 25*).

Nucleotide-free, rigor-like crystal structures reported for myosin and unconventional myosin V revealed distortional movements of the central β-sheet that rearrange the motor actin-binding region and reduce motor affinity for nucleotide prior to the power stroke (*26, 27*). We proposed previously that, like myosin, the central β-sheet of the kinesins undergoes structural changes involved in ADP release and force generation (*28*). We identified an invariant residue of the kinesin motor domain in a loop of the central β-sheet that could couple microtubule binding and release to ATP hydrolysis (*29*). Mutants of kinesin-14 Ncd that affect this residue, Y485, bind more tightly to microtubules than wild-type, hydrolyze ATP faster, and move faster in gliding assays (*29*). The most severe mutant, NcdY485K, results in the assembly of elongated oocyte spindles in vivo, evidence for its gain-of-function effects in live cells.

We undertook structural studies on the kinesin-14 NcdY485K mutant under the supposition that nucleotide and microtubule binding interactions by the mutant might favor previously unobserved transition states, causing them to be more highly populated without otherwise changing the motor mechanochemical cycle relative to wild-type kinesin. We observe features that have not been reported for previous kinesin structures, including a fluctuating power stroke, a post-hydrolysis state bound to ADP + free Pi, and shortening of β-strands prior to Pi release and the recovery stroke. Our new findings explain the coupling between ATP binding and motor interactions with microtubules, which underlie force production by the motor, and the basis of the enhanced mechanical output by NcdY485K compared to Ncd. The finding that the kinesin-14 working stroke fluctuates between the pre- and post-power stroke reveals an indeterminate mechanism of force generation by the motor upon ATP binding with multiple metastable states.

### MT-Ncd cryo-EM structures in three nucleotide states

We performed structural analysis on a truncated form of the Ncd motor (hereafter, denoted NcdY485K or Ncd) that contains the coiled-coil stalk, which dimerizes the motor and is essential for Ncd minus end-directed motility (*30, 31, 14, 21*), together with the Y485K mutation (*29*) (Fig. 1A). The tighter microtubule binding by NcdY485K is advantageous for solving high-resolution cryo-EM structures of the motor-microtubule complex. We solved the cryo-EM structures of microtubule-bound Ncd (MT-Ncd) in three different nucleotide states: i) NF state, ii) ATP state, and iii) ADP·Pi transition state (Table 1; Fig. 1, B-D and fig. S1). We applied MiRP2 (*32*) for high-resolution analysis of 13- and 14-protofilament (PF) pseudo-helical microtubules. We obtained models for Ncd in the NF, ATP, and ADP·Pi states bound to 13 PF and 14 PF MTs at 3.24 to 4.02 Å resolution (fig. S1 and fig. S2, table S1). Because the 14 PF maps were of higher quality, we used the 14 PF maps for the structural analysis reported here.

**Fig. 1.**
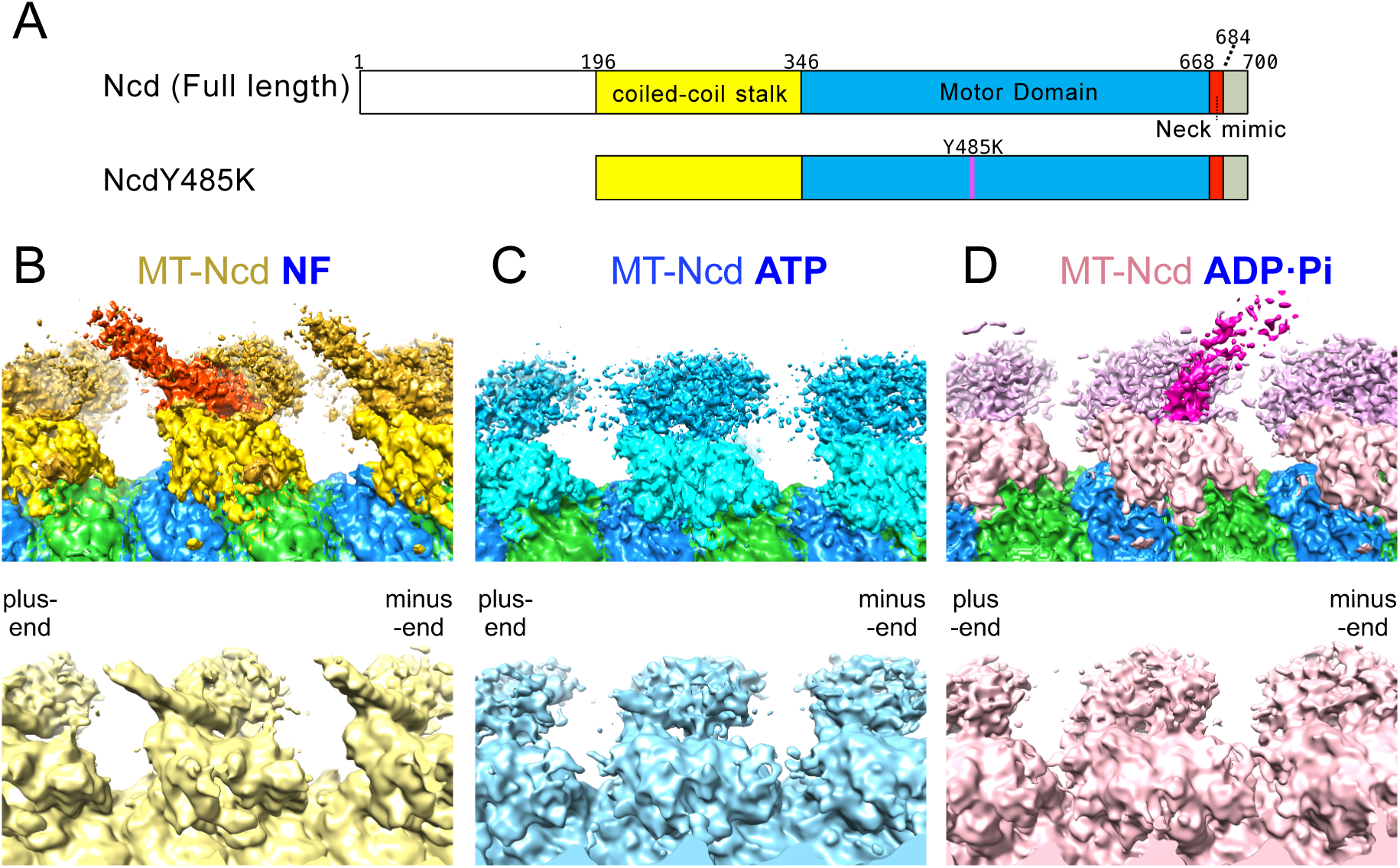
Kinesin-14 Ncd domains and MT-Ncd cryo-EM nucleotide states. (**A**) Ncd domain architecture. Ncd (Full length) and NcdY485K (mutated residue, magenta) analyzed in this study. The Ncd neck mimic (red) consists of residues 669 to 684 (16 residues). Stalk, yellow; motor domain, blue. (**B-D**) Cryo-EM density maps of NcdY485K bound to a microtubule in three nucleotide states. Top, ɑ-tubulin is green and β-tubulin, blue. Bottom, 6 Å lowpass filtered maps. The microtubule plus end is on the left and the minus end is on the right in each image. (**B**) MT-Ncd NF state. Top, the MT-bound NcdY485K head is yellow, the unbound head is dark gold, and the coiled-coil stalk is orange. (**C**) MT-Ncd ATP state. Top, the MT-bound NcdY485K head is cyan and the unbound head is dark cyan. (**D**) MT-Ncd ADP·Pi strong binding state. Top, the MT-bound NcdY485K head is pink, the unbound head is purple, and the stalk is magenta.

**Table 1.**
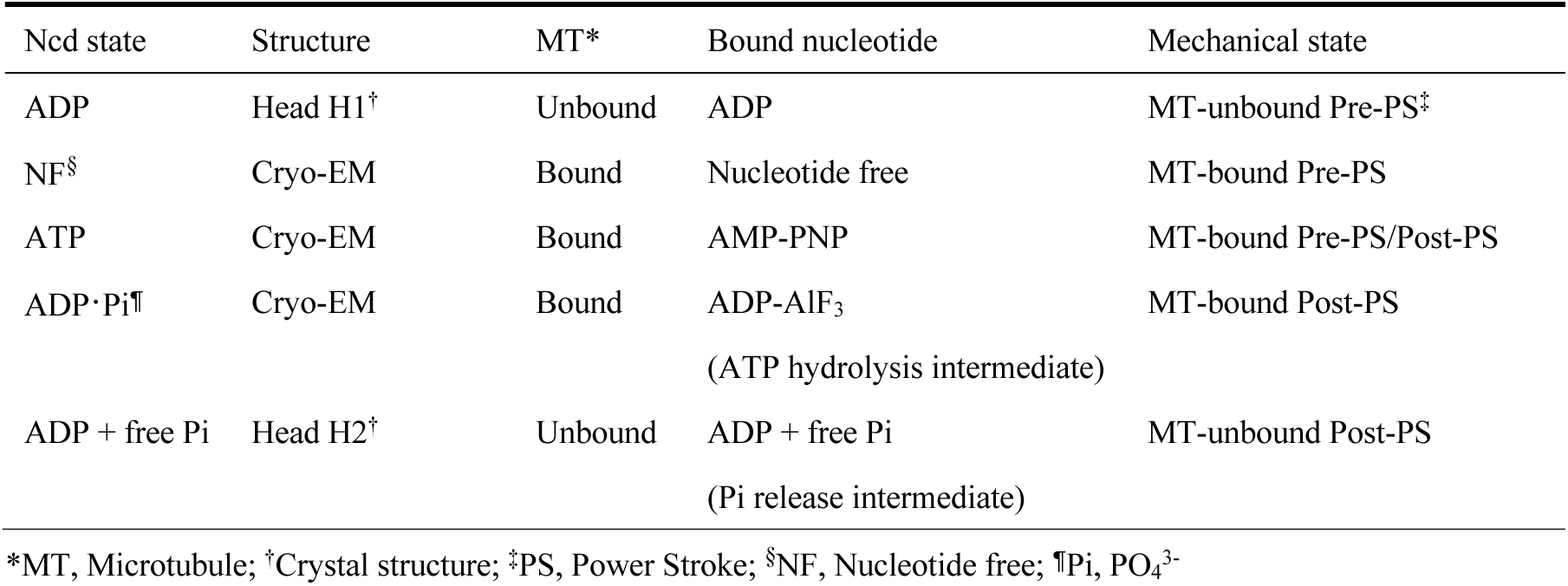
New kinesin-14 Ncd structures: nucleotide and mechanical states.

We could clearly see density in each state corresponding to the MT-bound and unbound Ncd motor domain, or head, together with density for the coiled-coil stalk (Fig. 1, B-D). Strikingly, the position of the stalk differed in each nucleotide state. The stalk was tilted towards the microtubule plus end in the NF state, it showed weak density that could be resolved into two conformations in the ATP state (see below), and it was tilted towards the microtubule minus end in the ADP·Pi state (Fig. 1, B-D). We interpret the stalk in the NF state to represent the MT-bound pre-PS conformation and the stalk in the ADP·Pi state to show the MT-bound post-PS conformation (Table 1). This interpretation is consistent with lower resolution MT-Ncd cryo-EM reconstructions (fig. S3) (*21*).

### MT-Ncd nucleotide-free state

We built an atomic model of MT-Ncd in the NF state by first fitting a previously reported crystal structure of wild-type Ncd (PDB 1CZ7, dimer 2, chains C and D) into the cryo-EM map. The crystal structure shows Ncd with both heads bound to ADP, unattached to microtubules (*33*). The 1CZ7 structure produced the initial MT-Ncd NF cryo-EM model. When fitting the motor domain of 1CZ7 chain C into the MT-bound head cryo-EM density, we found a slight tilt of the helical stalk (Fig. 2 and fig. S4). To further fit the stalk and unbound head, we applied a 6 Å lowpass filter to the map to enhance the signal-to-noise ratio of the stalk and unbound head densities. The 1CZ7 chain D could then be rigidly fitted into the cryo-EM density corresponding to the unbound head without the tilt of the stalk, resulting in an unambiguous fit of the crystal structure into the density corresponding to the stalk and unbound head. The model was then further refined (Fig. 2).

**Fig. 2.**
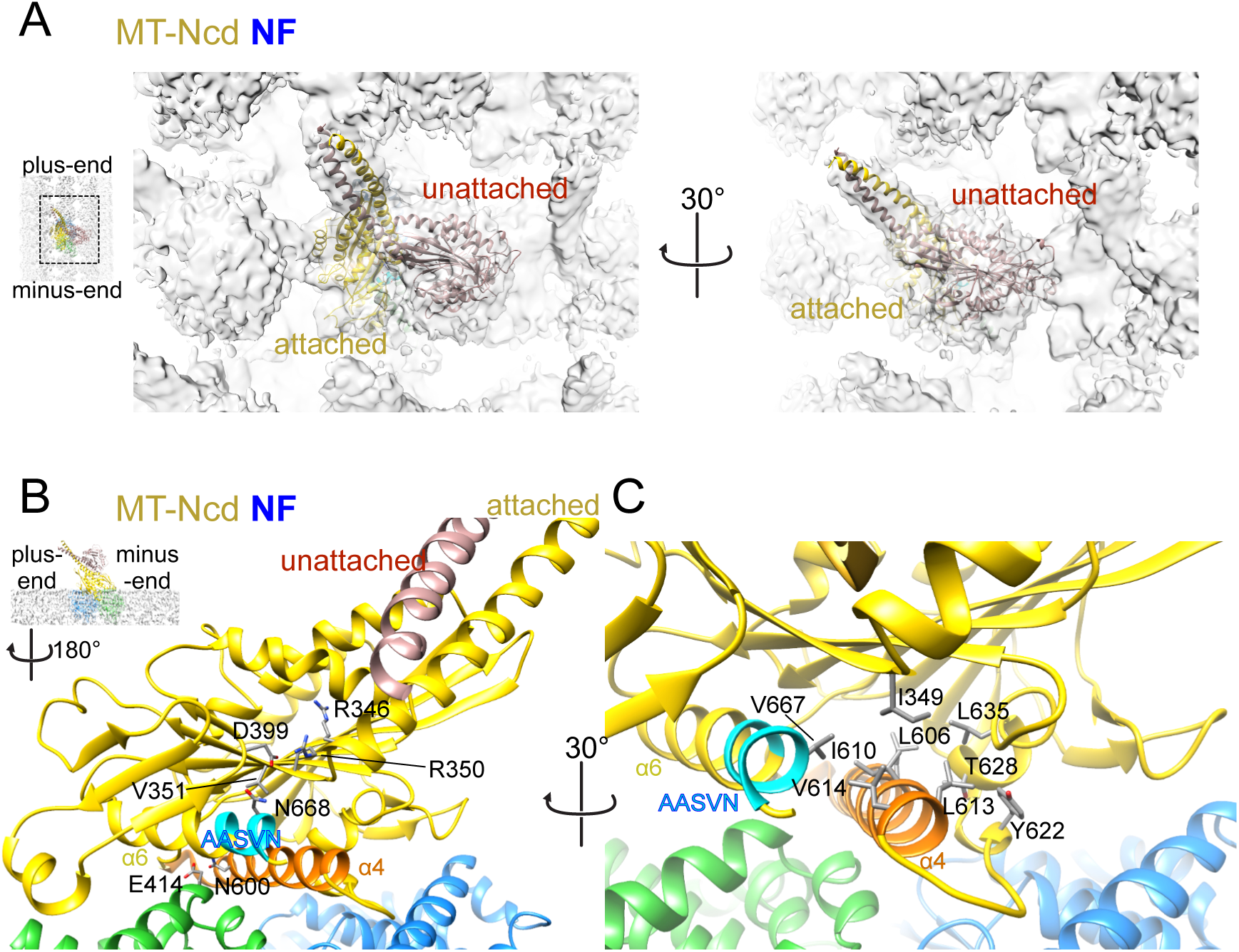
MT-Ncd nucleotide-free state. (**A**) MT-Ncd NF cryo-EM structure. Cryo-EM density was filtered by a 6 Å lowpass Gaussian filter. The MT-attached head is yellow, and the unattached head is light brown. (**B, C**) Cryo-EM model of the NcdY485K attached head in the NF state. The AASVN motif at the C-terminal end of helix α6 and base of the neck mimic, which is required for Ncd minus-end motility (*20*, *21*), is colored cyan, switch II helix ɑ4 is orange, ɑ-tubulin is green, and β-tubulin, blue. (**B**) Hydrophilic or (**C**) hydrophobic residue interactions near switch II helix ɑ4 and the AASVN motif, and hydrophobic side chains around the base of the stalk (stick models).

NcdY485K in the MT-Ncd NF model closely resembles the wild-type Ncd 1CZ7 structure (*33*) (dimer 2, chains C and D; RMSD of NF bound head and chain C head for 310 paired residues from N348 to N667 is 1.090 Å) (fig. S4A). After alignment of the motors by the NF unbound head or 1CZ7 chain D head, the MT-Ncd NF bound head is displaced by ∼7° towards the microtubule plus end compared to the 1CZ7 chain C head. By contrast, the NF unbound head and stalk are oriented the same as the crystal structure chain D (fig. S4, A and B), implying the presence of ADP in the unbound head. The Ncd neck mimic – the C-terminal region of the kinesin-14 motors that is structurally homologous to the neck linker of kinesin-1 and other plus-end directed kinesin motors (*24*) – is disordered in the MT-bound Ncd NF cryo-EM structure and is not built into the model. Movements are also observed in the switch I/II motifs, which undergo large changes in conformation during the nucleotide hydrolysis cycle. As observed in plus-end directed kinesins (*34, 35*), microtubule binding induces a loop-to-helix transition of switch II helix α4 at its N terminus. This conformational change results in the formation of an ion:dipole interaction (*36*) between N600 of Ncd helix ɑ4 and E414 of ɑ-tubulin helix H12 (Fig. 2B), which forms the initial binding interaction between the motor and tubulin (supplementary text). Binding by Ncd N600 to tubulin causes a mutant that converts the residue to lysine to have a strong dominant effect on microtubule binding (*37*). The NcdN600K mutant binds 2-3x more tightly to microtubules than wild type and blocks activation of the motor ATPase by microtubules (*37*) – the strong ionic interactions between the mutated NcdN600K residue and ɑ-tubulin E414 trap the motor in a nucleotide-free motor-microtubule complex. Microtubule binding also stabilizes the kinesin-14 Ncd helix ɑ6 C-terminus at the base of the neck mimic containing the AASVN motif (residues 664-668; fig. S5), which is required for Ncd minus-end directed motility (*22, 23*). The AASVN motif forms a hydrophobic interface that includes strand β1, loop L12, and helix ɑ4 and ɑ5 residues (Fig. 2C). N668 is close to D399, which forms a hydrophilic interaction with V351 of strand β1, stabilizing the stalk at its base (Fig. 2B). These interactions induce a small (∼3°) tilt of the stalk, reducing the distance between loop L10 and the stalk (fig. S4B).

### MT-Ncd ADP·Pi state

We reconstituted the ADP·Pi state by incubating Ncd with ADP-AlF_3_ attached to microtubules and analyzed by high-resolution cryo-EM. An atomic model of MT-Ncd in the ADP·Pi state was built by first fitting chain B of Ncd PDB 1N6M, in which the stalk bound to head H1 of chain A is rotated towards the microtubule minus end and thought to mimic the rotated stalk after ATP binding (*14–17*), into the cryo-EM density of the MT-bound head. The structure of the bound head differs from Ncd PDB 1CZ7 (*33*). It was manually refined using a crystal structure of human kinesin-1 in the ADP·Pi state bound to ɑ,β-tubulin (PDB 4HNA) (*35*) and an AlphaFold2-predicted structure (*38*) of full-length monomeric Ncd (AF-A0A0B4LI25-F1-model_v4.pdb) as guides (fig. S6A). A 6 Å resolution lowpass filtered electron density map was used to model the unbound head and stalk. We first rigidly fit the Ncd 1N6M chain A motor domain and stalk into the density of the unbound head. We then tilted the stalk ∼35°, measured from the bound head, to fit into the density and refined the model, which fit unambiguously into the density (fig. S6, A and B).

A striking feature of the MT-Ncd ADP·Pi structure is that the stalk has undergone a large rotation from its position in the MT-bound NF structure (Fig. 1, B-D and movie S1) and is tilted toward the MT minus end, stably positioned in the MT-bound post-PS conformation. The stalk angle of MT-Ncd ADP·Pi structure differs from Ncd crystal structures (fig. S6C, see below) (*14, 33*). When aligned by the unbound head, the MT-bound head is tilted by ∼10° more perpendicular to the microtubule surface (fig. S6, C and D), compared to Ncd 1N6M, which is interpreted to represent an unbound post-PS state (*14*).

Another striking feature of the MT-Ncd ADP·Pi cryo-EM structure is that the neck mimic at the C terminus of the motor is fully visible (Fig. 3 and fig. S6A). In the MT-Ncd ADP·Pi transition state cryo-EM structure, rotation of switch II helix ɑ4 at the motor-microtubule binding interface causes Ncd ɑ4 N600 to move away from ɑ-tubulin H12 E414, disrupting their interactions (Fig. 3A, supplementary text). In addition, the stalk rotation from the pre- to post-PS conformation alters AASVN interactions so that N668 is now close to R350, but retains its hydrophilic interactions with D399 in the loop between strands β1c and β2. N348, at the base of the stalk, has turned to face D399 and N668 and may form hydrophilic interaction through water, stabilizing the rotated stalk.

**Fig. 3.**
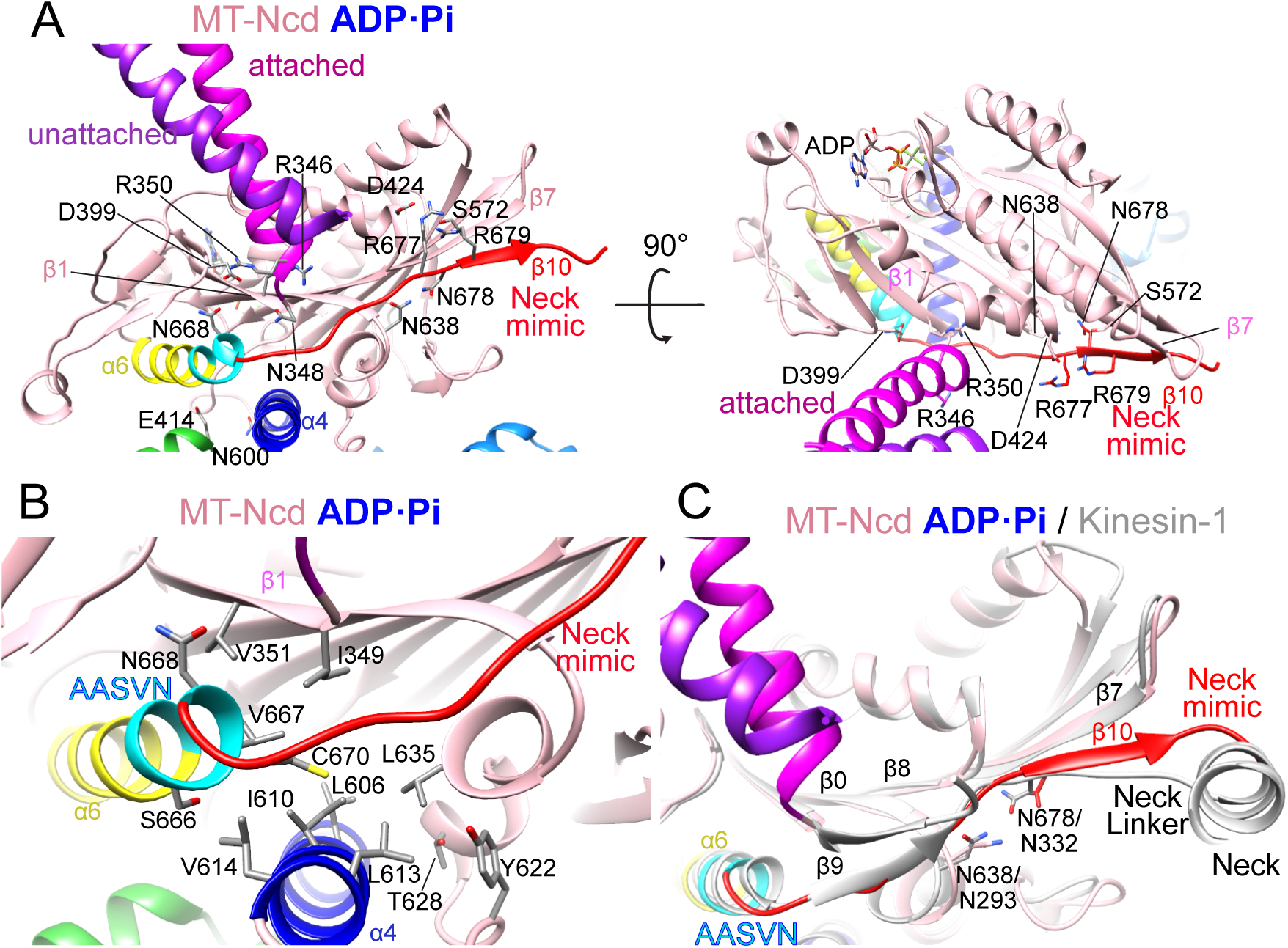
MT-Ncd ADP·Pi state. (**A**) Cryo-EM structure of MT-Ncd (motor domain, pink) bound to an ADP·Pi transition state analogue (*39*). The stalk helix of the MT-attached head is magenta, and the stalk helix of the unattached head is purple. Ncd switch II helix α4 is dark blue, helix α6 is yellow with the AASVN motif at the C-terminus of helix α6 shown in cyan, and the neck mimic is red. α-Tubulin, green; β-tubulin, blue. (**B**) Hydrophobic residues around helix α4, the AASVN motif, and the base of the stalk at the motor-MT interface. (**C**) Superposition of the Ncd ADP·Pi (pink) and kinesin-1 (KIF5B) PDB 1MKJ (gray) motor domain showing the kinesin-1 two-stranded cover neck bundle (strands β0 and β9). Ncd neck mimic N638 and N678 align with kinesin-1 neck linker N293 and N332, as shown.

The rotated stalk in the MT-Ncd ADP·Pi state is stabilized by interactions of hydrophobic patches in strand β1, helix ɑ4, and helix ɑ6 with the neck mimic, which is docked onto the motor core along its length (Fig. 3, A and B). As in the NF state, helix ɑ6 AASVN motif residues stabilize the base of the neck mimic by hydrophobic contacts with strand β1 and helix α4 residues (Fig. 3B). Neck mimic residues R677, N678, and R679 (fig. S5) interact with or are positioned close to helix α3 D424, loop L13 N638 and strand β7 S572, respectively, at the edge of the central β-sheet, stabilizing interactions with the motor core (Fig. 3A). The MT-Ncd ADP·Pi cryo-EM structure compared with a plus-end directed kinesin-1 crystal structure (PDB 1MKJ) shows that the docked neck mimic overlaps with the neck linker in position, but has different interactions with the motor core (Fig 3C). A major difference is that the cover strand, β0, at the N-terminus of the kinesin-1 motor domain forms a small two-stranded β-sheet, the cover-neck bundle, with strand β9 of the neck linker (Fig. 3C). Kinesin-14 lacks the cover strand and the cover-neck bundle, thus docking of the neck mimic onto the motor core is likely to produce less force than docking of the neck linker, although the initial region of the neck mimic stabilizes the stalk at its base by forming a shallow hydrophobic pocket (Fig. 3, B and C). Moreover, β10 of both the Ncd neck mimic and kinesin-1 neck linker form H-bonds to β7 of the central β-sheet, extending the β-sheet (Fig. 3C and movie S2) and altering its structure (table S2). Further, residues in the middle of the Ncd neck mimic (N678) and kinesin-1 neck linker (N332 in 1MKJ) both interact with loop L13 of the LGGN motif (N638 in Ncd or N293 in kinesin-1; Fig. 3, A and C), which is conserved in the kinesin superfamily proteins (*40*).

We also analyzed the changes in Ncd electrostatic potential upon neck mimic docking. Before neck mimic docking, the surface of the region involved in docking is negatively charged (fig. S6E, red dashed circle). After docking, the neck mimic covers the negatively charged surface and converts it to a positive charge (fig. S6F, blue dashed circle). Charge repulsion and steric clashes of side chain residues between the stalk and the neck mimic would drive the stalk from the pre-PS position, inducing stalk rotation to the post-PS position. The MT-Ncd ADP·Pi structure shows not only that the neck mimic resembles the docked neck linker of plus-end directed kinesin-1 (*11, 41*) but that it induces rotation of the stalk and is docked onto the edge of the central β-sheet following the power stroke, distorting the β-sheet (table S2).

### MT-Ncd ATP state

For the MT-Ncd ATP cryo-EM map, no model exists, including from previously reported crystal structures. We initially fit our MT-Ncd NF and ADP·Pi cryo-EM models into the electron density (Fig. 4 and fig. S7). For the density corresponding to the attached head, the MT-Ncd NF and ADP·Pi bound heads appeared at first glance to fit. However, careful examination of the MT-Ncd ATP density map revealed that structural elements were mismatched in the fits with both models (fig. S7). Ambiguities in the ATP cryo-EM map were also evident from the finding of two regions of density corresponding to the base of the stalk (Fig. 4 and fig. S7). This was attributed to conformations of the MT-bound head in two or more metastable states, including the pre- and post-PS states. To classify these states, we applied 3D-focused classification, but the attempt failed. This suggests that the Ncd ATP state fluctuates between multiple indistinguishable states. The finding of two regions of density for the base of the stalk in the MT-Ncd ATP cryo-EM map implies that the stalk fluctuates between the pre-PS and post-PS positions when the MT-bound head binds to ATP (Fig. 4A). This interpretation is consistent with lower resolution cryo-EM structures of Ncd bound to the ATP analogue, AMP-PNP, in which density attributed to the stalk is weak or not visible, compared to Ncd with no nucleotide or bound to ADP·Pi (*21, 42*).

**Fig. 4.**
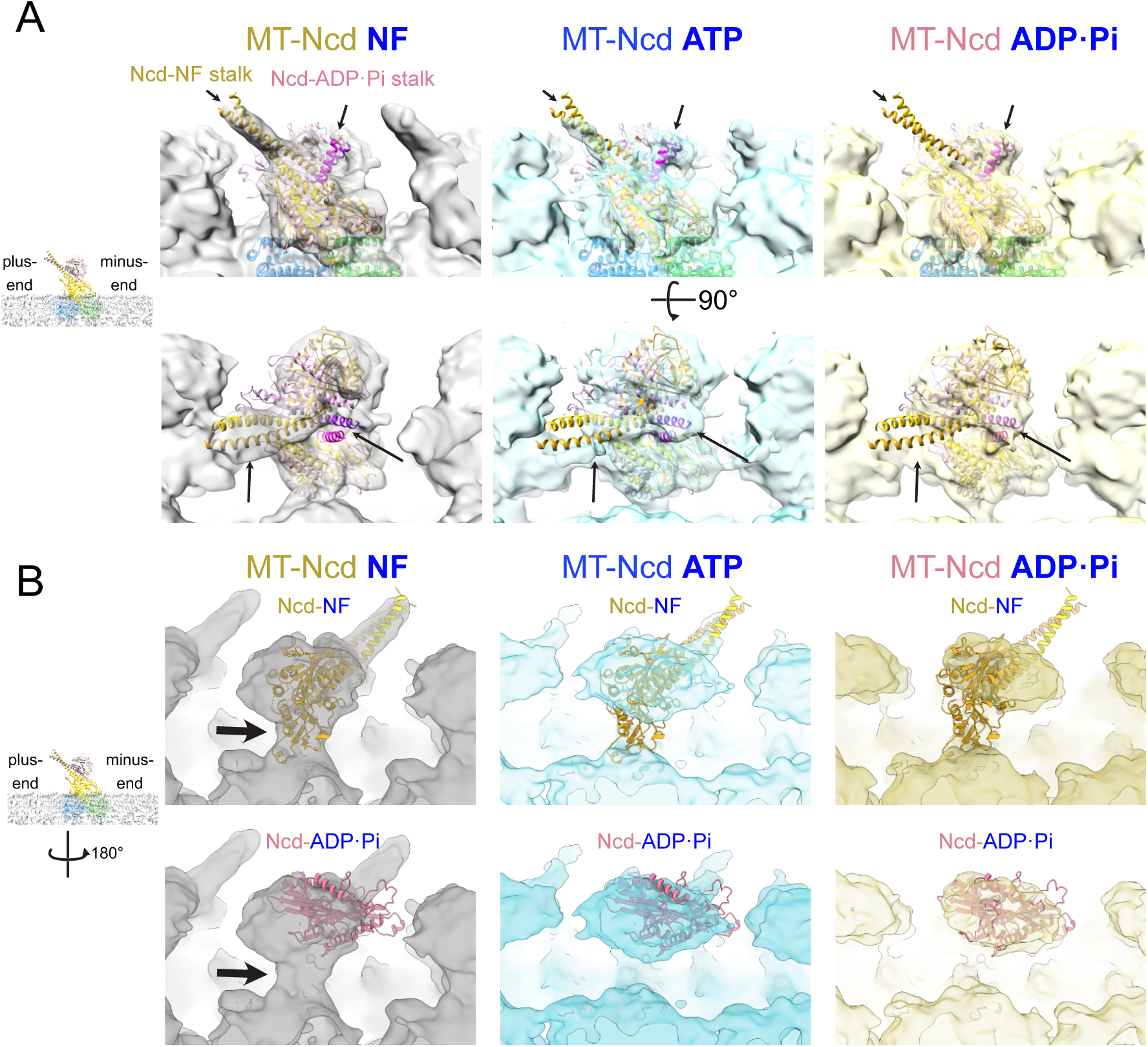
MT-Ncd ATP state. (**A)** MT-Ncd NF (left, gray), ATP (middle, cyan) and ADP·Pi (right, yellow) cryo-EM density maps fit with both the MT-Ncd NF (dark yellow) and ADP·Pi (pink with shorter purple stalk) cryo-EM models. A 10 Å lowpass Gaussian filter was applied to the maps. Top, side view; bottom, top view. Ncd stalk, black arrows. Inset (top), lower magnification MT-Ncd NF model showing the stalk tilted towards the MT plus end. α-Tubulin, green; β-tubulin, blue. (**B**) Cryo-EM density corresponding to the MT-Ncd NF (left, gray), ATP (middle, cyan), and ADP·Pi (right, yellow) unbound head. Top, the Ncd NF (yellow) or bottom, ADP·Pi (pink) cryo-EM model was fit into the density by rigid body fitting of tubulins and the bound head. A 10 Å lowpass Gaussian filter was applied to the maps. Inset (left), lower magnification MT-Ncd NF model showing the rotation of the models for the views shown on the right.

A comparison of the cryo-EM density corresponding to the unbound head of the MT-Ncd ATP, MT-Ncd NF, and ADP·Pi structures is also consistent with fluctuation between pre- and post-PS states by the MT-Ncd ATP bound head. The density corresponding to the unbound head of the MT-Ncd ATP structure is not in the same position as the unbound head of the MT-Ncd NF or ADP·Pi models (Fig. 4, A and B). The MT-Ncd NF cryo-EM map shows density between the unbound head and the microtubule surface (Fig. 4B, arrows), which is not observed in the MT-Ncd ATP cryo-EM map; rotation of the unbound NF head is also required to fill the density. The shape of the density for the unbound head in the ATP state is similar to that in the ADP·Pi state, but the unbound head density in the ATP state is displaced towards the MT plus end compared to the ADP·Pi state. These results imply that the MT-Ncd ATP bound head fluctuates between multiple metastable states that are similar to the more stable MT-Ncd NF and ADP·Pi pre- and post-PS states, respectively.

### NcdY485K crystal structure bound to free Pi with docked neck mimic

The NcdY485K dimeric motor was crystallized in the presence of excess Pi, and the structure was solved by molecular replacement (table S3). The 3.1 Å resolution model shows new structural features of the Ncd motor following ATP hydrolysis and motor detachment from the microtubule. The NcdY485K heads are oriented differently from one another in the crystal structure (fig. S8A), indicating that they represent different states of the mechanochemical cycle. Head H1 resembles previous crystal structures of wild-type Ncd bound to ADP (e.g., PDB 1CZ7), which show extensive interactions of the two heads with the end of the stalk. In the NcdY485K crystal structure, the stalk retains its interactions with head H1 but has undergone a large rotation of ∼80°, reorienting the stalk and head H1 relative to head H2 (fig. S8A), as in other Ncd wild-type and mutant crystal structures (*14–17*). Both heads contain density corresponding to ADP bound to the active site, but the head H2 active site also contains density for Pi adjacent to the bound nucleotide (Fig. 5A and fig. S8A). Density for Mg^2+^, which is ∼4-fold smaller in mass than Pi, could not be identified with certainty. Strikingly, the neck mimic is visible in head H2 (Fig. 5, A and B), unlike previously reported Ncd crystal structures, where the neck mimic is disordered and not visible (*14–17*). We interpret head H1 of the NcdY485K crystal structure to represent a MT-unattached pre-PS state and head H2 to reveal a previously unreported MT-detached post-PS state (Fig. 5, C and D; supplementary text).

**Fig. 5.**
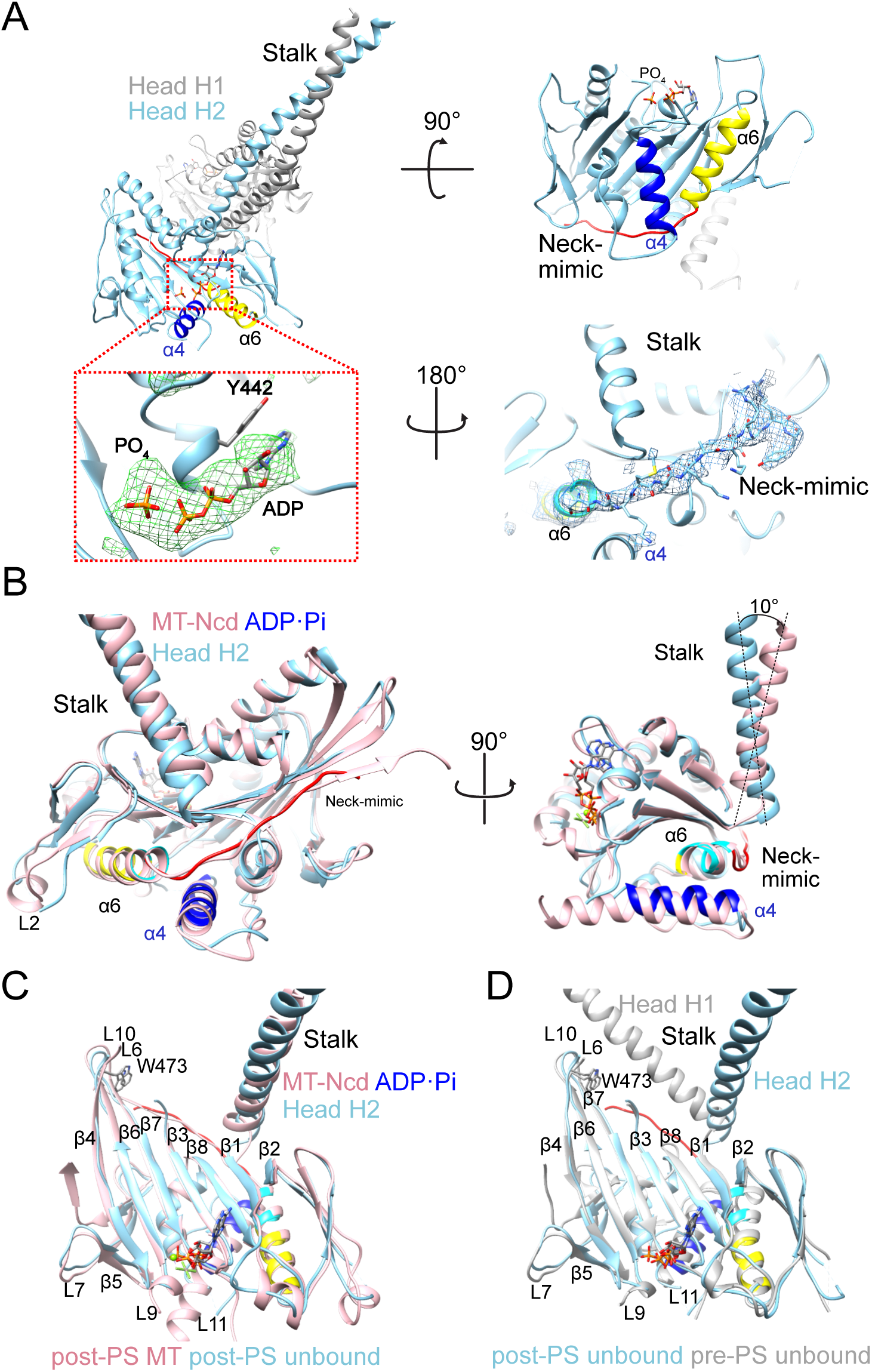
Ncd crystal structure in a MT-unattached post-PS state. (**A**) NcdY485K crystallized in the presence of high Pi concentration. Head H1 (chain A) is gray, and head H2 (chain B) is light blue. Head H2 switch II helix ɑ4 is dark blue and helix ɑ6 is yellow with AASVN shown in cyan. Inset, head H2 active site with Fo-Fc difference map (green mesh) contoured to 2.5 α and ADP + free Pi built into the density (stick models). Top (right), view from head H2 MT-binding surface. Neck mimic, red. Bottom (right), 2Fo-Fc difference map (blue mesh) showing density for head H2 neck mimic (stick models) contoured to 0.8 Λ. (**B**) Ncd head H2 (light blue) helix α4, helix α6, neck mimic and stalk compared with those of MT-Ncd ADP·Pi bound head (pink). (**C**) Ncd head H2 (light blue) superposed with MT-Ncd ADP·Pi bound head (pink; movie S2) or (**D**) Ncd head H1 (white; movie S3).

Several differences exist between head H2 of the crystal structure and the bound head of the MT-Ncd ADP·Pi cryo-EM structure, which represent different post-PS states. First, switch II helix α4 in head H2 of the crystal structure has undergone a helix-to-loop transition, shortening the structured region of the helix (L603-L614, 12 aa) by ∼50% at both its N- and C-terminus relative to the MT-Ncd ADP·Pi bound head (M593-Q614, 22 aa) (Fig. 5B) and MT-unbound kinesin motors, including plus-end directed kinesins (*43, 44*). However, the tilt of helix α4 in head H2 of the crystal structure is similar to that of the MT-Ncd ADP·Pi bound head (Fig. 5B), implying that the nucleotide state of head H2 is close to that of the MT-bound motor after microtubule detachment. The neck mimic in head H2 of the crystal structure (S669-Y680, 12 of 16 neck mimic residues) is visible and docked onto the motor core. The head H2 neck mimic and docked neck mimic in the MT-Ncd ADP·Pi bound head (C670-S684) differ in length and also by the inclusion of a residue, S669, that is present in helix α6 of the MT-Ncd ADP·Pi bound head. This residue has unwound to form the first residue of the neck mimic in head H2 of the crystal structure. Ncd neck mimic strand β10, which forms hydrogen bonds with the end of β7 in the ADP·Pi bound head, is unstructured or disordered in head H2 of the NcdY485K crystal structure (Fig. 5C). Further, the neck-mimic in head H2 of the crystal structure has a high B-factor compared to other regions of the motor that include head H1 and the stalk (fig. S8B). The structural elements at the head H2 motor-microtubule binding interface also show high B-factors, including strand β4 and/or loop L7 – the site of the NcdY485K mutation – and L11. Nonetheless, these regions are well defined in the electron density map (fig. S8B). The high B-factors of these structural elements indicate increased flexibility accompanying motor detachment from the microtubule.

### NcdY485K central β-sheet twisting and strand-to-loop transitions

The NcdY485K crystal structure also shows large changes in the central β-sheet of head H2, compared to head H1 (fig. S9A, table S2 and movie S3). The central β-sheet of the kinesin motors is evolutionarily conserved with myosin (*5*). In myosin, twisting or distortional movements of the β-sheet have been proposed to close the major 50 kDa cleft, driving ADP release and the swing of the lever arm during the power stroke (*26, 27*). The myosin β-strands that undergo the largest distortional changes between nucleotide states are strands 5-7, which precede the switch II loop, follow switch I, or are adjacent to switch I. Myosin strands 5-7 correspond to kinesin β7, β6, and β4, respectively (*28*). Residues in these three strands of NcdY485K head H2 show large changes in side chain position and orientation compared to head H1 (fig. S9, D and E). The movements are especially pronounced at the ends of strand β4 and involve W473 at the N terminus of β4 or present in the adjacent loop L6, depending on the nucleotide state (Fig. 5, C and D, and fig. S8, C-E), and the Y485K residue at the C terminus of β4 (or in loop L7) (fig. S9, D and E)) near the motor-microtubule binding interface. W473 is close to loop L10, which interacts with the stalk in the unbound and MT-bound pre-PS states (fig. S8D). The β-sheet twisting is first observed in the MT-Ncd NF state (table S2, fig. S9B and movie S4) and is increased in the MT-Ncd ADP·Pi bound head following the power stroke (table S2, fig. S9C, movie S1 and movie S5), and in Ncd head H2 after release from the MT (table S2), implicating β-sheet distortional changes in triggering stalk movements in kinesin-14 that produce force and displacements along microtubules.

In addition to the twisting of strands in the central β-sheet, extensive changes in the lengths of strands β4, β6, and β7 are observed between head H1 and H2 of the NcdY485K crystal structure (fig. S9A) that are caused by the conversion of the β-strands in head H2 into loops. Differences in β-strand length are also observed between head H1 and H2 of previous Ncd crystal structures (table S4), as well as between head H2 of the crystal structure and the MT-Ncd NF or ADP·Pi bound head, both of which have longer β-strands than head H2 (fig. S9, B and C). The differences in β-strand length between head H1 and H2 of the previous Ncd crystal structures could be due to differences in the post-PS transition states that the structures represent, some of which may correspond to states after Pi release. The strand-to-loop transitions in NcdY485K head H2 reduce the lengths of β4, β6, and β7 by as much as ∼40-70%, compared to head H1 or the MT-Ncd NF and ADP·Pi bound heads. The NcdY485K head H2 central β-sheet has higher B-factors (Cα, 99.6 to 167.8 Å^2^) compared to head H1 (Cα, 65.5 to 120.5 Å^2^) (fig. S8B), evidence of increased flexibility of the head H2 central β-sheet. We hypothesized that the head H2 strand-to-loop transitions are induced by ADP + free Pi bound to the head H2 active site after ATP hydrolysis that causes structural changes associated with motor detachment from the microtubule.

We tested this hypothesis by adding excess Pi to NcdY485K, which is assumed to be bound to 1:1 molar amounts of ADP after purification (*45*), and measuring changes in NcdY485K intrinsic fluorescence produced by tryptophan (Trp) residues to detect β-sheet conformational changes. The NcdY485K dimer construct that we analyzed (Fig. 1A) contains two Trp residues in each chain, both present in strands of the central β-sheet: W370 in strand β1a is pointed inward and protected from solvent in NcdY485K heads H1 and H2, whereas W473 is buried in head H1 under the ends of strands β6 and β7, and adjacent L6 and L10 residues (Fig. 6A). By contrast, W473 in head H2 is exposed to solvent by the strand-to-loop transitions of β6 and β7, and of β4 adjacent to W473 (Fig. 6A). Fluorimeter assays showed overlapping excitation spectra for NcdY485K (Ex_Max_, 275±4 nm, mean±SD, n=4) and free Trp (274±1 nm, n=3; P=0.9850, unpaired t test), but emission spectra showed a shift by ∼25 nm to shorter wavelength by ∼25 nm for NcdY485K (Em_Max_, 333±3 nm, n=14) compared to free Trp (357±2 nm, n=20; P<0.0001) (Fig. 6B and fig. S10, A and B). The blue-shifted NcdY485K emission spectrum is consistent with greater protection of the Trp residues from solvent in the motor relative to free Trp (*46*). Moreover, the addition of excess Pi to the NcdY485K motor, but not to free Trp, caused a significant reduction in fluorescence (Fig. 6C), consistent with greater solvent exposure of the Trp residues in the motor with Pi. Despite the reduction in fluorescence, the added Pi did not cause a shift in the NcdY485K Em_Max_ spectrum (fig. S10, A and B). The absence of a similar fluorescence decrease following addition of Pi to free Trp indicates that the effect is specific to NcdY485K.

**Fig. 6.**
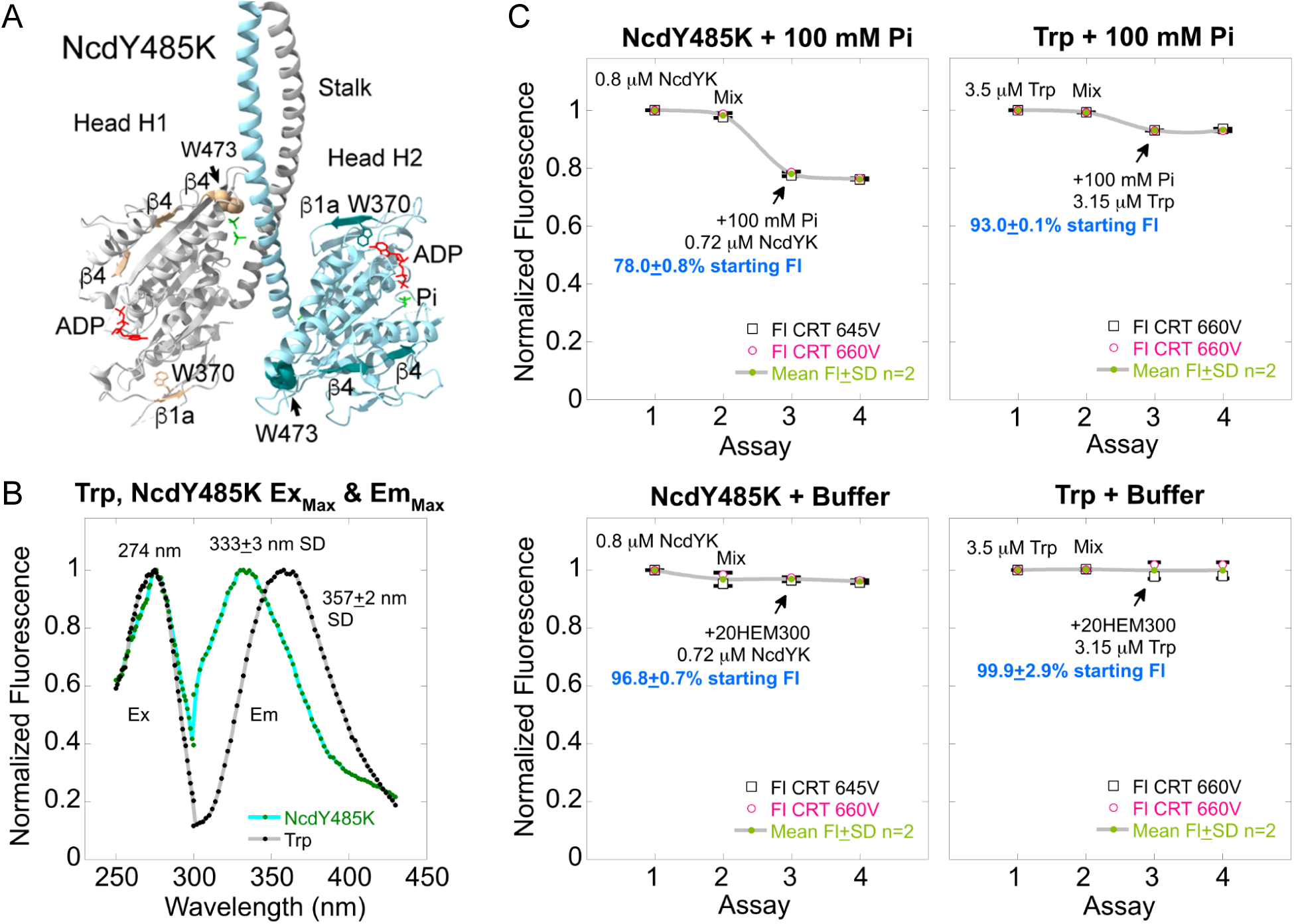
NcdY485K intrinsic fluorescence assays to detect central β-sheet conformational changes. (**A**) NcdY485K residues W370 (yellow, stick model) and W473 (dark yellow, space filled) in head H1 (white), and W370 (cyan, stick model) and W473 (dark cyan, space filled) in head H2. (**B**) Free tryptophan (Trp, gray) and NcdY485K intrinsic fluorescence (cyan) excitation and emission spectra. The NcdY485K emission spectrum is blue-shifted by ∼25 nm relative to the free Trp spectrum, indicating that the NcdY485K Trp residues are relatively buried. Ex, Excitation; Em, emission; Max, Maximum. (**C**) NcdY485K fluorescence assays. Top, NcdY485K (left) or free Trp (right) assays, consisting of an initial fluorescence assay, followed by an assay to control for mixing effects and two assays after adding 100 mM NaPi (Na3PO4). Bottom, NcdY485K (left) or free Trp (right) assays in which the same volume of buffer was added instead of NaPi. Plots show the mean±SD for assays performed under identical conditions of slit width, averaging time, and assay time. Data for NcdY485K are from paired buffer and control assays performed at the same PMT voltage (V), which was close to or the same as the free Trp assays. Free Trp assays shown were performed at the same PMT V. Fl, fluorescence. CRT, all assays were corrected for dilution due to addition of NaPi or buffer, and for photobleaching (see fig. S10C).

Without excess Pi, the active site of both NcdY485K heads is expected to be bound to ADP, like head H1 of the crystal structure, with the overall NcdY485K structure resembling a wild-type Ncd ADP-bound pre-PS structure (e.g., PDB 1CZ7). Upon binding to Pi, head H2 is expected to undergo structural changes resembling the NcdY485K post-PS crystal structure, including β-strand to loops transitions that expose W473 to solvent, which is predicted to decrease motor intrinsic fluorescence (*46*). We interpret the decrease in NcdY485K fluorescence observed in the fluorimeter assays upon addition of Pi to be caused by Pi binding to the active site of head H2, inducing β-strand transitions to loops in the head H2 central β-sheet. A potential effect of the strand-to-loop conversions, together with the central β-sheet strand twisting, is the storage of free energy to drive Pi release and the motor recovery stroke.

## Discussion

### Kinesin power stroke and recovery stroke

We report new structures that show five different MT-bound and unbound states of a kinesin motor before, during and after the power stroke (Table 1). Note that the ADP state (NcdY485K crystal structure head H1) is the same conformation as a previously reported wild-type Ncd motor bound to ADP, free in solution (*33*), and we could not build a MT-Ncd ATP atomic model because of the fluctuations between different states. Despite these caveats, the new structures reveal previously unreported features of the conformational changes by a kinesin motor during the mechanochemical cycle that are relevant to the motor energy transduction mechanism (Fig. 7, movie S6).

**Fig. 7.**
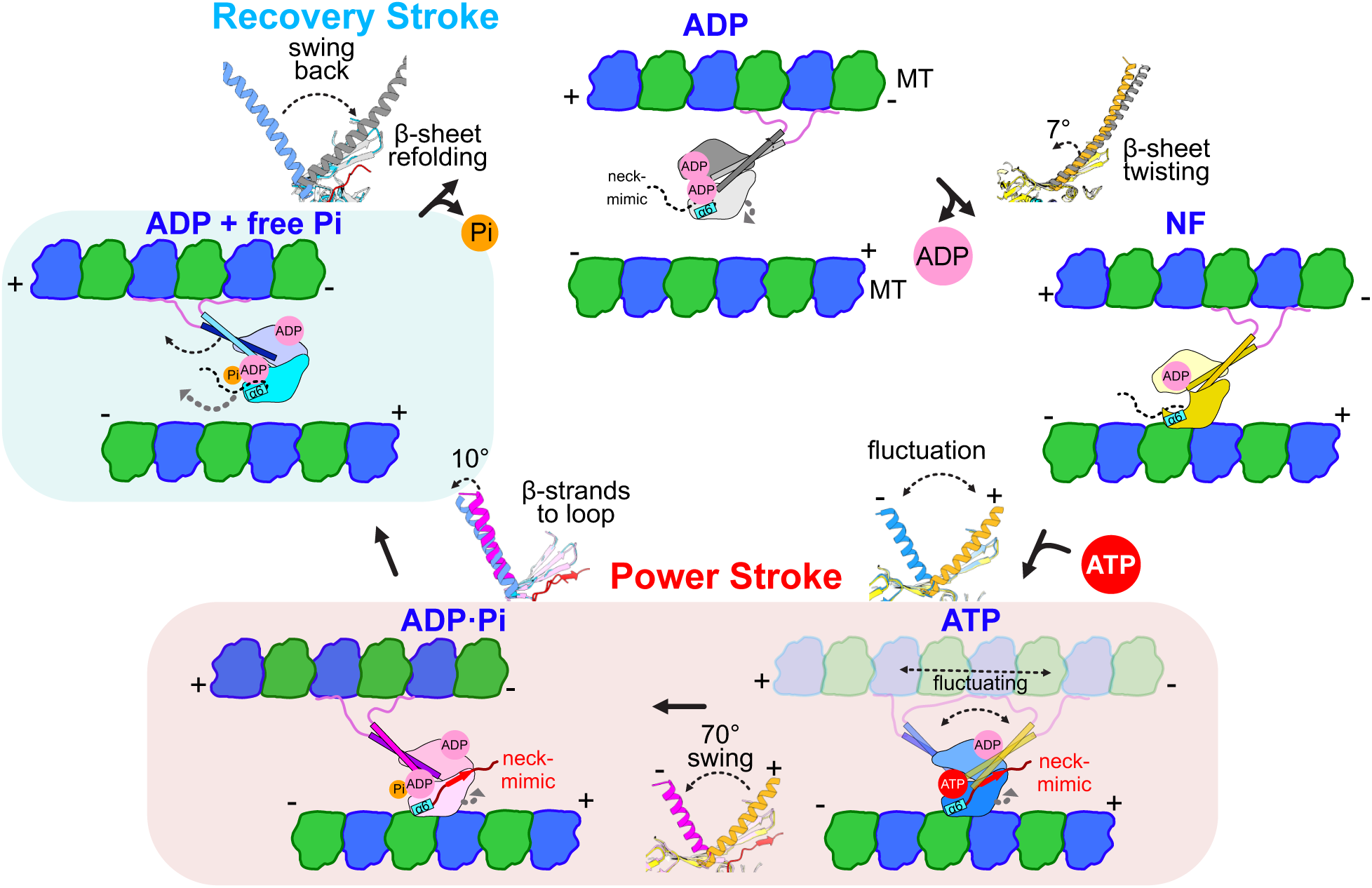
Model for kinesin-14 motility. Steps of ATP binding and hydrolysis for kinesin-14 minus-end microtubule movement: i) ADP state (unbound), ii) NF state (MT-bound), iii) ATP state (fluctuating, MT-bound), iv) ADP·Pi state (MT-bound), v) ADP + free Pi state (unbound) (Table 1). Following Pi release, the motor undergoes a recovery stroke to return to the ADP state (movie S6). Stalk helix (ribbon diagrams) angle changes are shown in each state.

Prior to microtubule binding, the kinesin motor, free in solution, has ADP bound to the active site (*45*) (Fig. 7, ADP). The motor binds to a microtubule by one head, releasing ADP from the bound head (*47*) (Fig. 7, NF). Helix ɑ6, which contains the AASVN motif required for kinesin minus-end motility (*22, 23*), interacts with β1 and helix ɑ4, tilting the stalk towards the microtubule plus end, stabilized by loop L6 and L10 interactions. The MT-bound head binds to ATP, causing the stalk to fluctuate in rotating towards the microtubule plus and minus ends (Fig. 7, ATP). After ATP binding and hydrolysis, the stalk is stably rotated towards the microtubule minus end (Fig. 7, ADP·Pi), consistent with a previous report of Ncd bound to an ADP·Pi analogue and microtubules by cryo-EM (*21*). Switch I is close to the position in the bound head that is occupied by γ-phosphate, and the neck mimic is docked onto the motor core at the edge of the central β-sheet. Neck mimic docking occurs with tilting of switch II helix α4 and the C-terminus of helix ɑ6, paralleling helix α4 and α6 movements upon docking of the kinesin-1 neck linker onto the motor core (*34, 35*). Detachment of the motor occurs with bound ADP + free Pi (*48*), the stalk rotated towards the microtubule minus end, and the neck mimic docked (Fig. 7, ADP + free Pi). Pi release, together with structural changes that untwist and refold the central β-sheet, return the MT-detached motor to the ADP form, repriming the motor for the next cycle of MT binding and nucleotide hydrolysis.

The NcdY485K motor reported here differs from wild-type kinesin-14 Ncd by interactions of both head H1 and H2 with switch I, which undergoes large changes in conformation during ATP hydrolysis. Y485K tilts towards and touches E483, causing the switch I loop, L9, to move close to the free Pi bound at the active site in H2 (fig. S11, supplementary text). These interactions with ATP prior to hydrolysis would orient and stabilize the bound nucleotide, explaining the faster ATP hydrolysis by NcdY485K compared to wild type without otherwise changing the motor mechanochemical cycle. Initiation of the switch I interactions by Y485K together with E483 would also account for the less severe effects of the NcdY485N and NcdY485E mutants on ATP hydrolysis (*29*).

### Kinesin-14 power stroke fluctuations

Fluctuations between the pre- and post-power stroke during ATP binding by NcdY485K observed in this study have not been reported previously (table S5, supplementary text). The fluctuations show that the helical stalk rotates alternately towards the microtubule plus and minus ends during the power stroke. The reversals in stalk rotation explain the plus-end directed displacements by a bidirectional kinesin-14 mutant, which were also observed for the wild-type motor and inferred to be due to movements of the stalk (*20*). The fluctuating stalk movements indicate that the kinesin-14 power stroke is indeterminate, relying on concerted movements of other structural elements, such as the neck mimic (see below) to produce a working stroke of the motor. The fluctuations could also explain the effects of an *Aspergillus* kinesin-14 microtubule-binding tail on motor directionality as constraining effects by the tail on stalk rotation conferred by certain motility assay geometries (*49*). Finally, the fluctuating power stroke provides an evolutionary link to the plus-end movement of kinesin-1 by demonstrating that the kinesin-14 stalk rotation can occur in either direction, directing the motor towards either the microtubule plus or minus end.

### Kinesin-14 neck mimic and force production

Docking of the kinesin-14 neck mimic along the motor core was reported previously in a plant kinesin, KCBP, where a truncated, monomeric motor construct bound to Mg^2+^-ADP was crystallized in a form interpreted to represent the motor in an ATP state (*24*). The KCBP helix ɑ6-neck mimic junction sequence more closely resembles the kinesin neck linker sequence than that of other kinesin-14 proteins (fig. S5), explaining its high affinity docking to the motor core. The KCBP motor lacked a stalk, so that movements of the stalk could not be observed. The findings we report here, including the Ncd stalk fluctuations during ATP binding and neck mimic docking in the MT-bound and detached post-PS states have not been reported previously – they extend previous observations for the role of the neck mimic in kinesin-14 motor proteins. In plus-end directed kinesin motors, the cover neck bundle (*13, 50*), a two-stranded β-sheet formed by cover strand β0 and neck linker strand β9, is thought to trigger neck linker docking, stabilized by interactions of a conserved neck linker residue (β10 N334 in kinesin-1) with β7 of the central β-sheet, producing the power stroke.

The absence of the cover strand and cover neck bundle in kinesin-14 likely affects the docking of the neck mimic and its stability in head H2 of the NcdY485K crystal structure. ATP hydrolysis and microtubule detachment cause changes in switch I/II, including a helix-to-loop transition at the N-terminus of helix ɑ4 and strand-to-loop transitions in the central β-sheet, described above. These structural changes probably weaken neck mimic interactions with the central β-sheet, destabilizing neck mimic docking in the Ncd ADP + free Pi state, which causes the stalk to rotate slightly towards the minus-end, but maintain its post-PS orientation.

### Kinesin structural changes during Pi release – central β-sheet unfolding & refolding

The formation of loops from β-strands and the high B-factors of the neck mimic, switch I/II and other structural elements at the motor-MT binding interface in the Ncd ADP + free Pi state (fig. S8B) indicate that these regions exhibit high entropy, which is likely to be associated with destabilization of the bound Pi and its release following motor detachment from the microtubule. The β4-L7-β5 cluster is close to the MT-binding surface in the MT-Ncd ADP·Pi bound head, and Y485K at the distal end of strand β4/loop L7 touches switch I. Switch I, in turn, is thought to interact with and sense the nucleotide phosphate bound to the active site (*51*). The end of strand β6, together with the other end of β4 and loop L10, interact with each other and with the coiled-coil stalk in the pre-PS state (fig. S8D). These strands and loops are coupled to one another by the central β-sheet, which is essential to the energy transduction mechanism – the β-strands twist and shorten, acting like a spring to store free energy to drive the working strokes of the motor. The strands and loops form a pathway for transmitting changes at the motor microtubule-binding surface to the nucleotide-binding cleft (*40*), triggering the power stroke, and then to loop L10 via the active site to drive the recovery stroke. The coupled structural changes at the microtubule-binding interface, active site, central β-sheet, neck mimic, and stalk during the kinesin-14 cycle lead to the power stroke upon motor binding to ATP, producing force and microtubule sliding, and then induce Pi release and the motor recovery stroke after ATP hydrolysis and detachment from the microtubule.

## Materials and methods

### Protein expression and purification

A plasmid for bacterial expression of NcdY485K (MGSM-H293-K700-His6) was constructed in pMW172 (*29*) using usual recombinant DNA methods. The NcdY485K protein contains 54 residues of the α-helical coiled-coil Ncd stalk joined to the conserved motor domain, but is deleted for the N-terminal tail and ∼96 residues of the predicted stalk. The motor domain contains a residue change of Y485K.

pMW172/NcdY485K was transformed into *E. coli* BL21(DE3) host cells (Novagen), cultured overnight, and grown in 1 L LB media in 2 L flasks (total 8 L) at 37°C in the presence of 100 μg/ml ampicillin to OD_600_ ∼0.6, then cooled down in cold water. Expression was induced by adding 0.1 mM isopropyl-α-D-thiogalactopyranoside (IPTG) overnight at 18°C. Cells were harvested and stored at -80°C until use. Cells were resuspended in buffer A [50 mM HEPES pH 7.5, 80 mM NaCl, 1 mM MgCl_2_, 1 mM EGTA, 1 mM DTT, 0.7 µM leupeptin, 2 µM pepstatin A, 1 mM phenylmethylsulfonyl fluoride (PMSF), and 2 mM benzamidine], lysed by sonication and cleared by centrifugation. The lysate was applied to a HiTrap SP ion exchange column (GE Healthcare), washed with 100 mM NaCl in HEM buffer (10 mM HEPES pH 7.5, 1 mM MgCl_2_, 1 mM EGTA, and 1 mM DTT), and eluted with a linear gradient of 500 mM NaCl in HEM buffer. The second peak of the eluate was concentrated and applied to a Superdex 200 Increase (GE Healthcare) size exclusion chromatography column equilibrated with 100 mM NaCl in HEM buffer for the final step in purification. Protein in the peak fractions was concentrated to ∼13-14 mg/ml, flash frozen in liquid nitrogen, and stored at -80°C.

NcdY485K protein for crystallization was expressed essentially as described above, but in *E. coli* Rosetta2 (DE3) pLysS host cells (Novagen). Cells were lysed by sonication in binding buffer (BB; 12 mM NaPO_4_ pH 7.4, 300 mM NaCl, 20 mM imidazole, and 0.5 mM MgCl_2_). After clearing by centrifugation, the lysate was loaded onto a Ni column for purification by Ni-affinity chromatography. Elution was with BB + 280 mM imidazole, followed by Superdex 200 size exclusion FPLC in HEM200 buffer (HEM + 200 mM NaCl). Purified NcdY485K protein was concentrated, frozen, and stored at -80°C.

Tubulin was purified from porcine brains purchased from a meat factory in Kobe, Japan, as described previously (*52*).

### Cryo-EM grid preparation

Tubulin (60 μM) was incubated in PEMGTP buffer (100 mM PIPES pH 6.8, 1 mM MgCl_2_, 1 mM EGTA, and 1 mM GTP) at 37°C for 30 min and microtubules were stabilized by adding 60 µM taxol. Ncd-microtubule complexes in different nucleotide states were prepared using essentially the same procedure except for the presence or absence of the nucleotide analogue in the Ncd protein solution. For nucleotide-free Ncd grid preparation, the purified protein was used. For AMP·PNP, 2 mM AMP·PNP was added to the purified Ncd protein and incubated for 10 min. For ADP-AlF_3_, 2 mM ADP, 2 mM AlF_3_, and 8 mM NaF were incubated with purified Ncd for 30 min. Microtubules (4 µl drop of 40 µM dilution) were placed onto a glow-discharged holey carbon grid (R2/1; Quantifoil) prepared on a JEC-3000FC (JEOL). After 30 s, the solution was wicked away with a piece of Whatman No. 1 filter paper, and 80 µM of Ncd protein in different nucleotide states was applied (4 µl drop), incubated for 1 min, blotted, and plunge-frozen in liquid ethane using a Vitrobot Mark IV (Thermo Fisher Scientific).

### Cryo-EM data collection and processing

Cryo-EM data were collected on a JEOL CRYO ARM 300 cryo-electron microscope at SPring-8 in Japan with a cold-field emission gun as the electron source, an in-column Ω filter, using a K3 direct electron detector (Gatan) in the electron counting mode. Images were collected at a nominal magnification of ×60,000, corresponding to a calibrated pixel size of 0.753 Å. Each movie was recorded in correlated-double sampling (CDS) mode for 2.5 s and subdivided into 50 frames with 50 e^−^/Å^2^ at the specimen. The image shift method automatically acquired the data using SerialEM software (*53*), with a defocus range of −0.8 to −1.5 μm. Approximately 6,000-8,000 movies were acquired for each condition, and the total number of images is described in table S1.

Collected movie frames were subjected to Relion implemented motion correction, and the contrast transfer function (CTF) of the movie frames was estimated by CTF in RELION 3.1 (*54*). To obtain high resolution maps of microtubules decorated by Ncd in different nucleotide states, we employed MiRPv2 (*32*), a method specialized for high resolution single particle analysis and for analyzing the pseudo-helical nature of microtubules with a seam, which can precisely determine the positions of α and β-tubulins and the seam. First, microtubules on the micrographs were manually picked using the helical mode of RELION (v3.1.4). The selected microtubules were then extracted as particles with a box size of 600 Å^2^ and box separation distance of 82 Å with 4 x binning. The extracted particles were 3D classified (PF-sorting) into 6 classes of 11- to 16-protofilaments (PF) using low pass filtered 11-to 16-PF microtubules as references. After PF-sorting, we reassigned incorrectly assigned particles in the same filament to the correct protofilament number, assuming that the protofilament number is the same throughout a single microtubule. Because most of the particles were classified as 13- and 14-PF (see fig. S1), we analyzed the predominant 13 and 14 PF classes. In the next step, we aligned the seam position. To improve the signal-to-noise ratio of the particles, neighboring particles in an MT were averaged over 7 segments along the helical axis and were used for 3D classification (*55*). The rotation angles were modified to ensure being linear in a microtubule followed the 3D classification. Moreover, X/Y shifts, which center the microtubule particles onto the reference, were also applied to ensure the linearity in a microtubule. To check the seam position, we 3D-classified the refined particles to the possible seam references (i.e., 26 references for a 13-protofilament microtubule, with 13 seam positions and their counterparts translated 1 monomer along the helical axis), and the seam position of particles was corrected according to the classified result. Finally, we reconstructed the particles by re-extracting particles from the micrographs without binning. The corrected particles were C1-reconstructed, and the refined particles were used in the symmetrized reconstruction to improve the resolution. The refined particles were further post-processed to acquire a high-resolution map according to the postprocess procedure of Relion. Figures and movies of cryo-EM structures were made in UCSF Chimera or ChimeraX (*56, 57*).

### Crystallization

Purified protein was thawed, incubated on ice with 5.3 mM Mg^2+^-ATP for 60 mins, then centrifuged for 6 min at 16,630x g and 4°C. Crystals were grown in hanging drops by mixing 1.3 µl protein + Mg^2+^-ATP with 1.5 µl reservoir solution, which consisted of 1 ml of 50 mM NaPO_4_ pH 6.8, 7 mM DTT, 10 mM MgCl_2_, 0.8 M NaCl, 13% w/v PEG8000 and 5% glycerol. Crystallization trays were incubated at 16°C. Crystals appeared in 2 days and were grown at 16°C for ∼6 days before harvesting. Crystals were incubated in cryoprotection buffer (reservoir solution + 25% glycerol), flash frozen, and stored in liquid nitrogen.

### X-ray diffraction data collection and processing

A complete diffraction data set was collected at the Advanced Photon Source Synchrotron on NECAT Beamline 24-ID-E with an Eiger detector (Argonne National Laboratory, Chicago, IL USA). The data set was processed using HKL2000 (*58*). The NcdY485K structure was solved by molecular replacement using the Phaser program (*59*) from the Phenix crystallographic suite. A wild-type Ncd structure (PDB 1CZ7) was used as the initial search model. PDB 1CZ7 contains two Ncd dimers in the asymmetric unit. Dimer 2, in which a longer α-helical coiled-coil stalk is visible (*33*), was used to solve the NcdY485K structure. Model building was performed using the Coot program (*60*), and refinement was carried out using Phenix Refine and REFMAC5 (*61, 62*). Refinement was continued until the R-values converged to 0.25 (R_free_=0.29) with 3.15 Å resolution. The model has good stereochemistry with all residues within the allowed regions of the Ramachandran plot, as analyzed by MolProbity (table S3) (*63*). Figures were made in UCSF Chimera or ChimeraX (*56, 57*).

### Fluorescence assays

NcdY485K intrinsic fluorescence due to tryptophan (Trp) residues was assayed to detect conformational changes that affect the Trp residue environment in different motor states. Assays were performed using FPLC-purified NcdY485K protein (*29*) at 0.36 to 2.71 µM in 20HEM300 buffer (20 mM HEPES pH 7.2, 1 mM EGTA, 1 mM MgCl_2_, 300 mM NaCl) and 22°C in a Cary Eclipse spectrofluorometer. Fluorescence was read before and after a mixing control (with no additions) and after adding 100 mM Na_3_PO_4_ pH 6.8. Free Trp at 0.72 to 13.22 µM was assayed as a control, together with controls performed by addition of the same volume of 20HEM300 buffer instead of Na_3_PO_4_. Fluorescence recorded during the assays was corrected for decreases in NcdY485K or free Trp concentration due to the increased volume upon addition of Na_3_PO_4_ or buffer, and for photobleaching using a free Trp photobleaching decay curve (fig. S10C). The rate of photobleaching was found to be 0.000517/s for the 20 nm slit width used under the conditions of the assays shown in Fig. 6C, which were performed for 10 s and 274 nm excitation with 358 nm emission for free Trp or 330 nm emission for NcdY485K; PMT voltages (V) were the same as matched control assays in which 20HEM300 buffer was added to the cuvette instead of 100 mM Na_3_PO_4_ pH 6.8. Fluorescence spectra (Fig. 6B and fig. S10A) were read using slit widths of either 10 or 20 nm, and were not corrected for photobleaching, as there was very little change in intensity in repeat assays.

The NcdY485K + Pi fluorescence decrease shown in Fig. 6C is interpreted to be caused by changes in the central β-sheet of head H2 that expose W473 to solvent. The free Trp + Pi fluorescence decrease is assumed to be due to Trp interactions with Pi. The small NcdY485K + buffer fluorescence decrease was greater in protein that had been freeze-thawed several times before assaying and is attributed to protein unfolding and loss upon mixing (e.g., by binding to the micropipette tip). Assays under different PMT voltage conditions or with a different purification of NcdY485K protein also showed a large decrease in fluorescence with added Pi (not shown; deposited in Zenodo as files with names that include the following terms: Fig. 6C, not shown, NcdYK prep 2). Protocols for the intrinsic fluorescence assays have been deposited into Zenodo with the data files.

### β-sheet structural analysis

A crystal structure of NcdY485K (this report), together with those of four other wild-type or mutant Ncd dimers with a rotated stalk was displayed in UCSF Chimera (*56*) and β-strand length was analyzed (table S4). The crystal structures show chain A (head H1) in a pre-PS state and chain B (head H2) in a post-PS state. A wild-type Ncd crystal structure with both heads in a pre-PS state was analyzed as a control (PDB 1CZ7 dimer 2, chains C and D). The Ncd dimer post-PS heads show shorter strands β7, β6, and β4, which correspond to β-strands 5-7, respectively, in nucleotide-free myosin – these strands in myosin undergo the largest changes between nucleotide states. The shorter head H2 β-strands are due to strand-to-loop transitions – the strands differ in length in different Ncd mutants, potentially reflecting different transition states of head H2 in the crystal structures (shortest β-strands, bold font in table S4). The two heads of previous Ncd dimer crystal structures have been reported to be bound to ADP. By contrast, NcdY485K head H1 is bound to ADP and head H2 to ADP + free Pi.

Analysis of central β-sheet residue movements between the NcdY485K crystal structure heads H1 (ADP) and H2 (ADP + free Pi) was performed by first identifying the residues that comprise each strand of the β-sheet. Residues were included in the analysis if they were present in a β-strand in at least one head of the motor and if the residue and atom numbers matched between the two heads. This allowed us to include the effects of β-strand to loop transitions, which are expected to increase the flexibility of the central β-sheet, and loop conversions to β-strands, which would decrease the β-sheet flexibility. As a result, strands were compared with loops if head H1 and H2 differed in β-sheet structure. The pre-stroke wild-type Ncd structure (1CZ7 dimer 2, chain C and D) was included as a control, but β-sheet residues were defined for the 1CZ7 heads separately from NcdY485K H1 and H2, although using the same criteria. This was necessary because the 1CZ7 structure represents a pre-power stroke state, differing from the post-stroke NcdY485K structure.

Differences in β-strand residue positions and orientations between the two heads of NcdY485K and those of Ncd 1CZ7 dimer 2 were calculated using the Chimera RMSD tool (*64*). The Chimera script used to analyze the residue differences is included in the Supplementary Materials. The RMSD output value for each NcdY485K β-strand residue is shown in a heat map (fig. S9E). The totals shown below the heat map were obtained by squaring the RMSD values, multiplying by the number of atoms in each residue of the β-strand, and summing the products. The β-sheet differences between the NcdY485K and Ncd 1CZ7 structures were compared by calculating the mean β-strand RMSD±SD from the Chimera RMSD output and determining the significance in GraphPad Prism using an unpaired t test (see fig. S9 legend).

The MT-Ncd NF (nucleotide-free) and MT-Ncd ADP·Pi (ADP-AlF_3_) cryo-EM structures (this report) were also analyzed for β-strand residue differences compared to the MT-unbound state that could be interpreted as central β-sheet twisting or distortional movements (table S2). The MT-Ncd NF and MT-Ncd ADP·Pi cryo-EM models are somewhat lower resolution (3.24-3.99 Å) than the NcdY485K crystal structure (3.15 Å), and the positions of side groups of the β-strand residues are expected to be less accurate in these structures. We therefore performed Cα-Cα RMSD analysis to provide an estimate of the changes in the β-sheet between the unbound Ncd ADP state and the MT-Ncd nucleotide states. This was done by aligning the NcdY485K crystal structure head H1 β-strand residue C_a_ atoms with the C_a_ atoms of the same residues of the MT-bound or unbound head of the cryo-EM models using Matchmaker in ChimeraX v1.6 (*57*). The RMSD values were displayed by Matchmaker after selecting the NcdY485K β-strands in the Sequence Viewer window and selecting *Also restrict to selection* for NcdY485K head H1 and *Iterate by pruning long atom pairs* in the Matchmaker window prior to running the analysis. Differences were observed for all the pairwise comparisons. The largest RMSD mean value was observed for the comparison of the NcdY485K crystal structure head H1 and H2 (table S2).

This method for estimating β-sheet RMSD values differs from that used for the NcdY485K crystal structure head H1 vs head H2 and shown in the heat map in fig. S9E. First, it does not include residues that are present in β-strands in the protein chain under comparison, but not in NcdY485K head H1. Second, Matchmaker uses only the single Cα atom per residue for fitting, rather than other atoms or all atoms of the residue. Third, Matchmaker uses only the paired Cα-Cα atoms to obtain an RMSD value. By contrast, the RMSD analysis shown in fig. S9E is an all-atom comparison of residues that are present in a β-strand in at least one head of the motor. The methods used to estimate β-sheet RMSD differences in table S2 are thus not the same as those used to produce the heat map in fig. S9E, since they use only a single point per residue, which is the Cα atom. Exclusion of side chain atoms is expected to result in under-estimates of the actual β-sheet differences. For example, the comparison of NcdY485K head H1 vs head H2 shown in table S2 gave a β-sheet Cα-Cα RMSD mean difference of 1.154 Å, compared to an all-atom RMSD mean difference of 1.250±0.230 Å (mean±SD, n=11 strands) for the analysis shown in fig. S9E (see legend). Because of the methods that were used for the analysis, the RMSD data in table S2 are not directly comparable to the data for the NcdY485K crystal structure shown in fig. S9E. However, the cryo-EM MT-bound head β-sheet distortional changes are apparent from the Cα-Cα RMSD mean values, and they are also observed in the superpositions of the MT-bound heads with NcdY485K ADP head H1 (fig. S9B and C) and movies showing morphs between each other, including stick models of the β-strand carbon backbones (movie S1), or between the MT-bound heads and H1 or H2 (movies S2-S4).

### Statistical analysis

Statistical tests were performed using GraphPad Prism software. Unpaired t tests were used to compare data for two independent groups. P values are given in the legends for the figures in which the data are shown.

## Acknowledgements

We thank G. Christoph, C. Gopalasingam, and K. Kato for cryo-EM data collection management and support at SPring-8 and K. Chin for research management support. S.A.E. thanks S. Zhang and J.J. Gooley for Duke-NUS NBD Programme laboratory space, D.M. Virshup for Duke-NUS CSCB Programme hospitality, T.G. Oas for guidance on the fluorescence assays, and E.C. Meng for information about Chimera Matchmaker. **Funding:** This work was supported by grants from the Japan Society for the Promotion of Science KAKENHI (22K06809 to TI, 22H02795 and 21H05254 to RN); AMED-CREST from the Japan Agency for Medical Research and Development (AMED; JP21gm1610003 to TI and JP21gm0810013 to RN); Japan Science and Technology Agency [JST/FOREST (JPMJFR214K to TI); JST/Moonshot (JPMJMS2024 to RN)]; Takeda Science Foundation and Mochida Memorial Foundation for Medical and Pharmaceutical Research to TI and RN; and the Uehara Memorial Foundation, Bristol-Myers Squibb, and Hyogo Science and Technology Association to RN. The cryo-EM experiments were performed at SPring-8 with the approval of the Japan Synchrotron Radiation Research Institute (JASRI) (Proposal No. 2021B2536, 2022A2536, 2022B2542, 2023A2542, 2023B2537). NcdY485K purification and crystallization were supported by Duke-NUS Medical School funds to SAE and NcdY485K structural analysis by NIH NICHD 1R21HD105034-01A1 Grant Award to SAE. JS acknowledges Ministry of Education National University of Singapore AcRF Tier-1 (R154-000-A72-114) and Tier-2 (R-154-000-B03-112) funding support. Diffraction data were collected at the Northeastern Collaborative Access Team beamlines, funded by NIH NIGMS P30 GM124165, using resources of the APS, a US DOE Office of Science User Facility operated for the DOE Office of Science by Argonne National Laboratory under Contract No. DE-AC02-06CH11357, using APS beamline 24-ID-E. The Eiger 16M detector on the 24-ID-E beam line is funded by NIH-ORIP HEI Grant S10OD021527. **Author contributions:** Conceptualization: SAE, TI, RN; Investigation: TI, HS, RN, SAE, YW, CJ; Data analysis: SS, TI, MYW, CJ, RN, SAE, HH, JS; Data curation: TI, SS, CJ, SAE, MYW; Funding acquisition: TI, RN, SAE, JS; Project administration: TI, SAE, RN, JS; Supervision: TI, SAE, RN, JS; Writing original draft: SAE, TI, RN; Figures and tables: TI, SS, MYW, SAE, RN; Review/editing: TI, SAE, MYW, RN, JS, CJ, YW; Final review: All authors. **Competing interests:** The authors declare that they have no competing interests. **Data and materials availability:** MT-NcdY485K cryo-EM density maps and atomic models have been deposited into the Electron Microscopy Data Bank (EMDB) and the Protein Data Bank (PDB) under the following accession codes: NF 13PF EMD-39664/PDB ID 8YY2, NF 14PF EMD- 39665/PDB ID 8YY3, AMPPNP 13PF EMD-39668, AMPPNP 14PF EMD-39669, ADP-AlF_3_ 13PF EMD-39666/PDB ID 8YY4, and ADP-AlF_3_ 14PF EMD-39667/PDB ID 8YY5. The cryo-EM raw micrographs will be deposited to EMPIAR. The NcdY485K atomic model coordinates and .mtz file containing the unprocessed x-ray data have been deposited into the PDB under accession code PDB ID 8YUE. The RMSD data shown in fig. S9E and the fluorimeter data shown in Fig. 6, B and C, and fig. S10 have been deposited into Zenodo (*65*).

## Supplementary Materials

Supplementary Text

Figs. S1 to S11

Tables S1 to S5

Computer Script

References 66-72

Movies S1-S6

## Notes

http://10.5281/zenodo.10853624

### Competing Interest Statement

The authors have declared no competing interest.

