## Supplementary_Material for "Structural transitions in kinesin minus-end directed microtubule motility"

**Affiliations:** [1]Division of Structural Medicine and Anatomy, Department of Physiology and Cell Biology, Kobe University Graduate School of Medicine, Kobe, 650-0017, Japan; [2]Department of Cell Biology, Duke University Medical Center, Durham, NC 27710 USA; [3]Structural Biology Division, Japan Synchrotron Radiation Research Institute, SPring-8, Sayo, Hyogo, 679-5184, Japan; [4]Neuroscience & Behavioral Disorders Programme, Duke-NUS School of Medicine, SG 169857; [5]Department of Biological Sciences, National University of Singapore, SG 117558

*Corresponding authors:


†Equal contribution

‡Current Address: Centre for Advanced Microscopy, The Australian National University, Canberra ACT 2601, AUS

**The PDF file includes:**

Supplementary Text
Figs. S1 to S11
Tables S1 to S5
Computer Script
References 66-72

**Other Supplementary Materials for this manuscript include the following:**

Movies S1 to S6

### Supplementary Text

#### Tubulin structural changes induced by NcdY485K binding

The MT-Ncd NF cryo-EM structure (this report) shows that motor binding to microtubules occurs with an initial interaction between Ncd N600 of helix α4 and E414 of α-tubulin helix H12. Binding by the Ncd motor appears to be stabilized by residues at the N terminus of helix α4 (R592-N600), which fit into a pocket formed by α-tubulin H11' and H12 (*28*). Binding by Ncd to tubulin in the nucleotide-free state causes a small displacement of H12 E414 towards the microtubule minus end, compared to the tubulin residue in an unbound microtubule (PDB 6DPV chain L**,** 3.3 Å MT-GDP, 14 protofilament). Ncd binding also causes α-tubulin helix H11 to move towards the microtubule plus end, inducing a conformational change of helix H9-loop S8 and a helix-to-loop transition of H6, which is shorter at its C terminus by three residues in the MT-Ncd NF cryo-EM structure than in unbound α-tubulin (PDB 6DPV).

The MT-Ncd ADP·Pi cryo-EM structure shows further changes in α-tubulin, including a rotation of E414 at the N terminus of α-tubulin helix H12, positioning it further away from NcdY485K N600. There are also changes in α-tubulin helix H11 that increase the helix length by three residues at the N terminus compared to α-tubulin of the MT-Ncd NF cryo-EM model. Taken together, the changes in α-tubulin between the MT-Ncd NF and ADP·Pi cryo-EM structures are relatively small (RMSD=0.520 Å, 427 pruned atom pairs; RMSD=0.529 Å, all 428 atom pairs). The changes observed in the MT-Ncd NF cryo-EM structure that occur upon motor binding to tubulin (RMSD=0.555 Å, 427 pruned atom pairs; RMSD=0.568 Å, all 428 atom pairs) are larger than those that occur in tubulin between the MT-Ncd NF and MT-Ncd ADP·Pi states. The initial changes that occur upon motor binding deform tubulin both at the motor-tubulin interface as well as further away from the site of motor binding to tubulin, reminiscent of those that are observed upon binding by kinesin-1 to microtubules (*66*). These deforming changes could have large effects on microtubule structure, which would not necessarily be detected in our studies, but could affect binding by additional motors, either Ncd or others, to the microtubule. If so, these effects could help explain the cooperativity of Ncd binding to microtubules (*42, 67*).

#### Ncd crystal structures after phosphate release

Four other Ncd crystal structures with one head in the pre-PS state and the other head in the post-PS state have been reported previously: wild-type Ncd (PDB 5W3D), NcdG347D (PDB 3U06), NcdT436S (PDB 3L1C) and NcdN600K (PDB 1N6M) (*14-17)* . When superimposed with MT-Ncd ADP·Pi (ADP-$AlF_3$; this report), the structures align well, but there are differences in residues in helices and strands, including rotation of N600 (or N600K in 1N6M) away from α-tubulin E414. Disruption of Ncd N600 or N600K interactions with α-tubulin indicates that the structures do not represent microtubule-bound forms of the motor, although they are close to the MT-Ncd ADP·Pi cryo-EM transition state. Instead, the Ncd crystal structures most likely represent unattached post-power stroke states of the motor that differ slightly from one another. The finding of residues at the proximal end of the neck mimic that are docked onto the motor core in two of the motors, NcdT436S (*15)* and NcdG347D (*16*), implies that these structures might be closer than the others to the MT-Ncd ADP·Pi state. However, the post-power stroke head H2 (in chain B) of both NcdG347D and NcdT436S shows a shortening of helix α4 at the N-terminus compared to the MT-Ncd ADP·Pi attached head, which is likely to accompany destabilization of microtubule binding and detachment by the motor from the microtubule.

#### NcdY485K mutant effects

The NcdY485K mutant alters the invariant EIY motif of the kinesin motor proteins. The new NcdY485K crystal structure shows that the EIY motif differs in conformation from previous wild-type and mutant Ncd crystal structures. The mutated Y485K residue in both heads H1 and H2 is tilted towards E483 and positioned close enough to E483 to form H-bonds (fig. S11A). By contrast, wild-type Ncd (PDB 5W3D; fig. S11, B and C) and other mutant Ncd crystal structures (1N6M, 3L1C, 3U06) show Y485 tilted away from E483 (n=4 structures, n=8 heads). Y485K in head H2 of the mutant has rotated towards E483, which enhances strand β4 twisting (fig. S11C). Head H2 β4 twisting positions Y485K and E483 near R539 of the switch I helix, α3, and R539 has moved close enough to the free Pi bound at the H2 active site to form hydrogen bonds (fig. S11A). The same interactions of H2 switch I R539 with ATP prior to hydrolysis would orient and stabilize the bound nucleotide, accounting for the increased NcdY485Y ATPase rate relative to wild-type Ncd (*29*).

**NcdY485K fluctuations during the ATP state**

For kinesin-14 NcdY485K, ATP binding occurs in multiple metastable states, leading to fluctuations during the power stroke (this report). This may be true for other kinesins (*12*). The power stroke of myosin and dynein is also thought to occur by a multistate process (*19, 69*) (table S5), resulting in tight filament binding and force production. Thus, the power stroke of all three cytoskeletal motors is characterized by multiple states, including the pre- and post-PS, that can be interpreted as representing metastable transitions. Multiple stable states are characteristic of a biological switch (*70*).

### Supplementary Figures

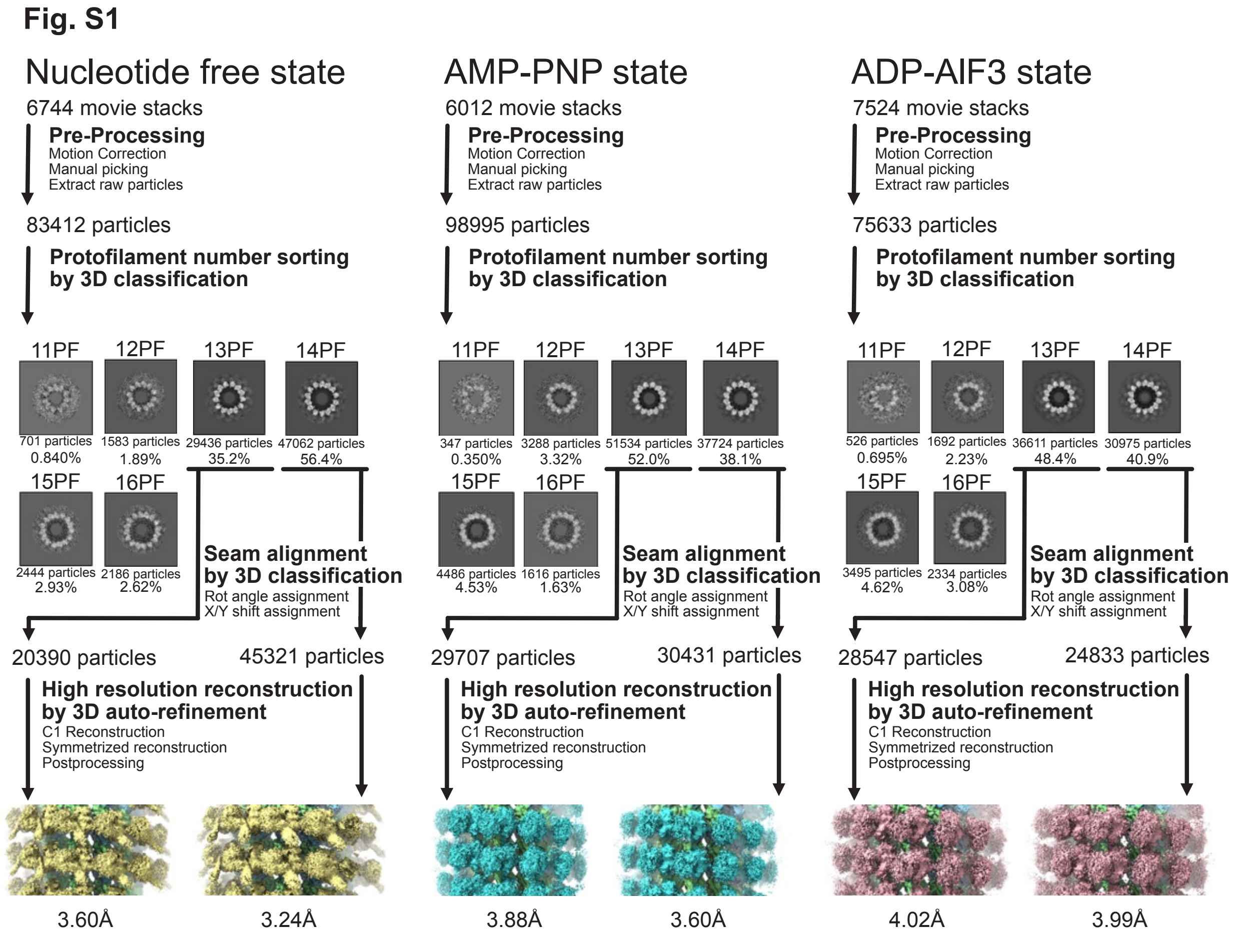


**Fig. S1. Workflow of cryo-EM 3D reconstruction of MT-Ncd complexes.** Flow chart of cryo-EM data processing and analysis. Helices were picked from micrographs and extracted as a particle with a box separation distance of 82 Å and 4 x binning. After sorting for protofilament (PF) number, seams were aligned, and 1 x binned particles were 3D reconstituted with symmetry.

**Fig. S2**

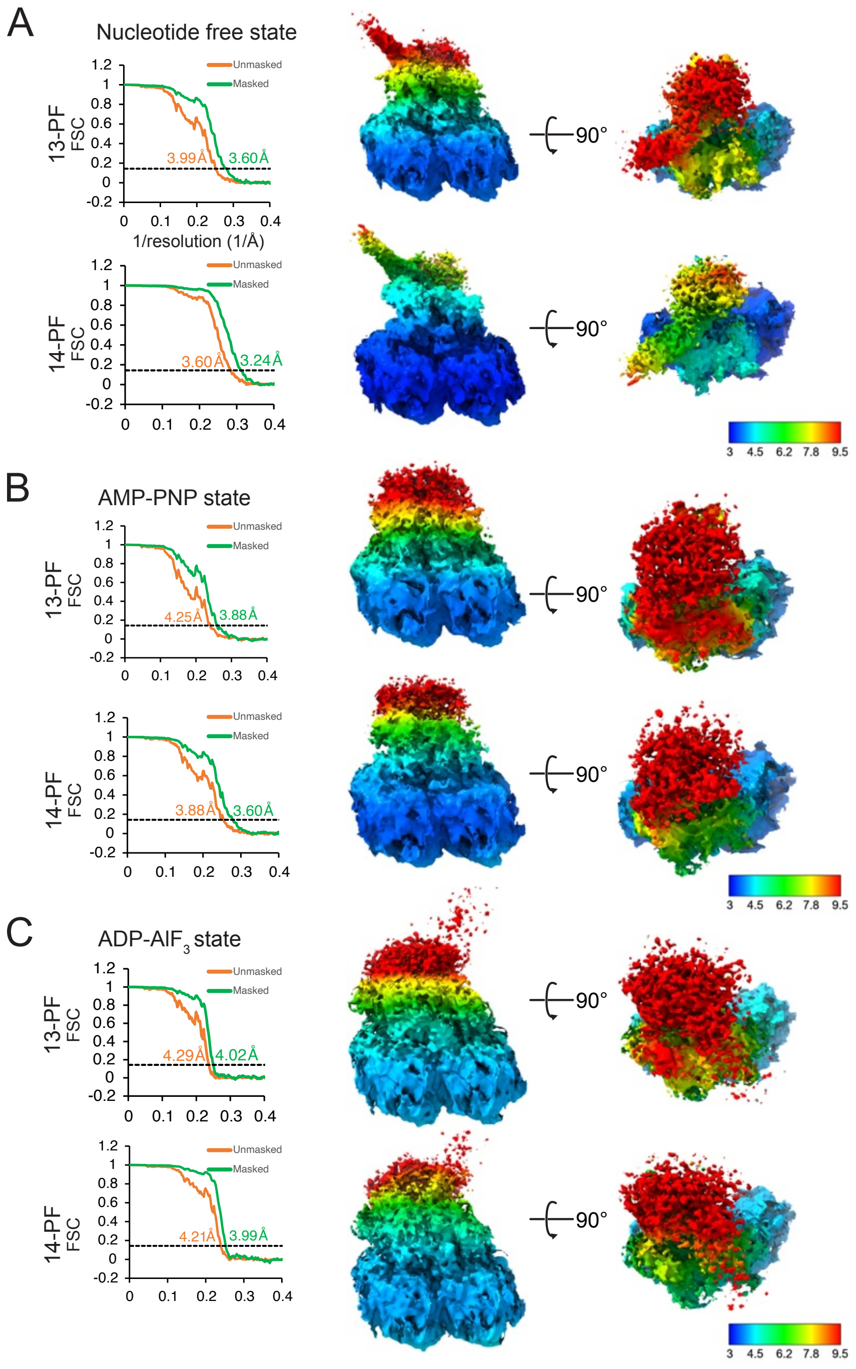


**Fig. S2. Cryo-EM map quality assessment.** (**A-C**) Fourier shell correlation (FSC) curves of the final cryo-EM maps of MT-bound Ncd in three different nucleotide states. Two views of the density maps are colored by resolution. Scale bars indicate the resolution corresponding to colors. (**A**) MT-Ncd NF (nucleotide-free) state, (**B**) MT-Ncd ATP (AMP·PNP-bound) state, and (**C**) MT-Ncd ADP·Pi (ADP-$AlF_3$ bound) state.

**Fig. S3**

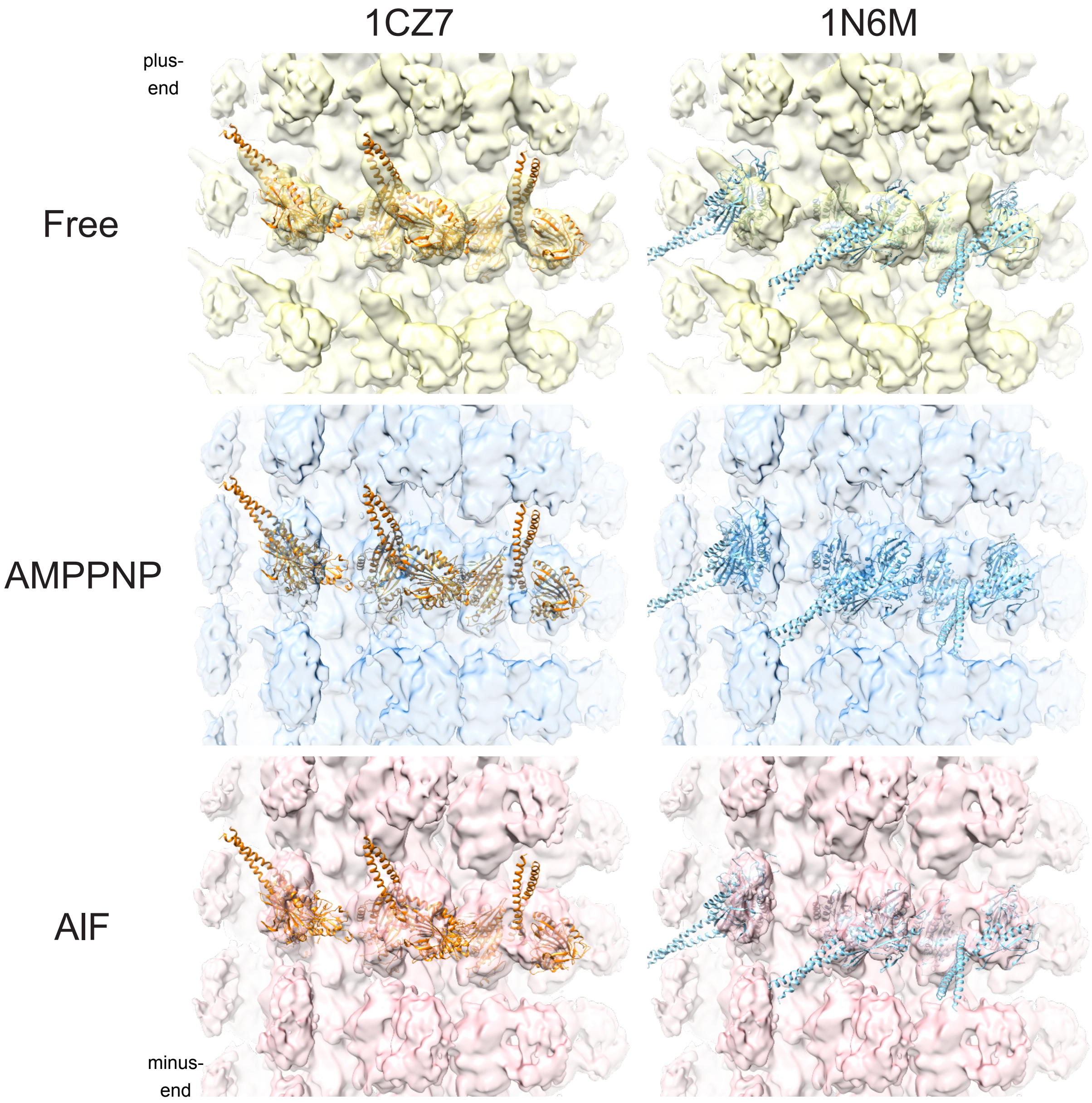


**Fig. S3. Overall structure of MT-Ncd complexes.** Ten angstrom low pass-filtered maps (top to bottom) superimposed with previously reported crystal structures (left, right). Top, MT-Ncd NF (Free, nucleotide-free); Middle, MT-Ncd ATP (AMPPNP); bottom, MT-Ncd ADP·Pi (AlF). Left, wild-type Ncd (PDB 1CZ7 dimer 2); right, NcdN600K (PDB 1N6M). 1CZ7 shows the Ncd motor-ADP with the unrotated stalk in a pre-power stroke (pre-PS) conformation, tilted toward the microtubule plus end. By contrast, 1N6M shows the Ncd motor-ADP with the stalk bound to head H1 rotated and tilted toward the microtubule minus end and head H2 in a post-power stroke (post-PS) conformation.

**Fig. S4**

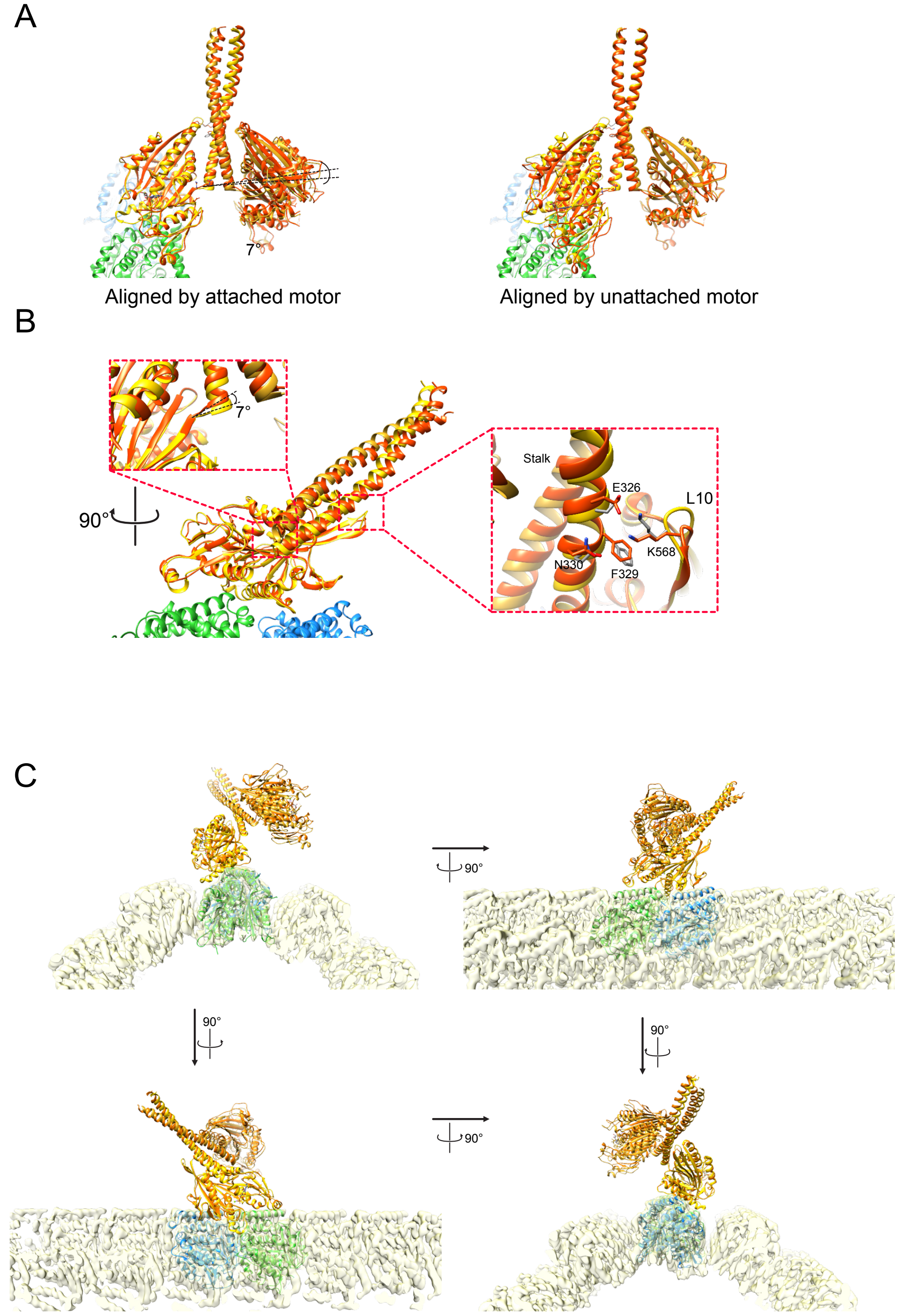


**Fig. S4. Structural comparison of MT-Ncd bound and unbound pre-PS state.** (**A**) Superposition of the MT-Ncd NF pre-PS cryo-EM structure (yellow) with an unbound Ncd ADP pre-PS crystal structure (PDB 1CZ7 dimer 2, chains C and D, orange), aligned by the MT-Ncd attached motor domain (left) or by the MT-Ncd unattached motor domain (right). (**B**) Insets, close-up views of (**A**) left, from different angles, showing the tilt of the stalk (top) and the stalk interaction with loop L10 (right). (**C**) Aligned structures in (**A**) left, viewed from different angles.

**Fig. S5**

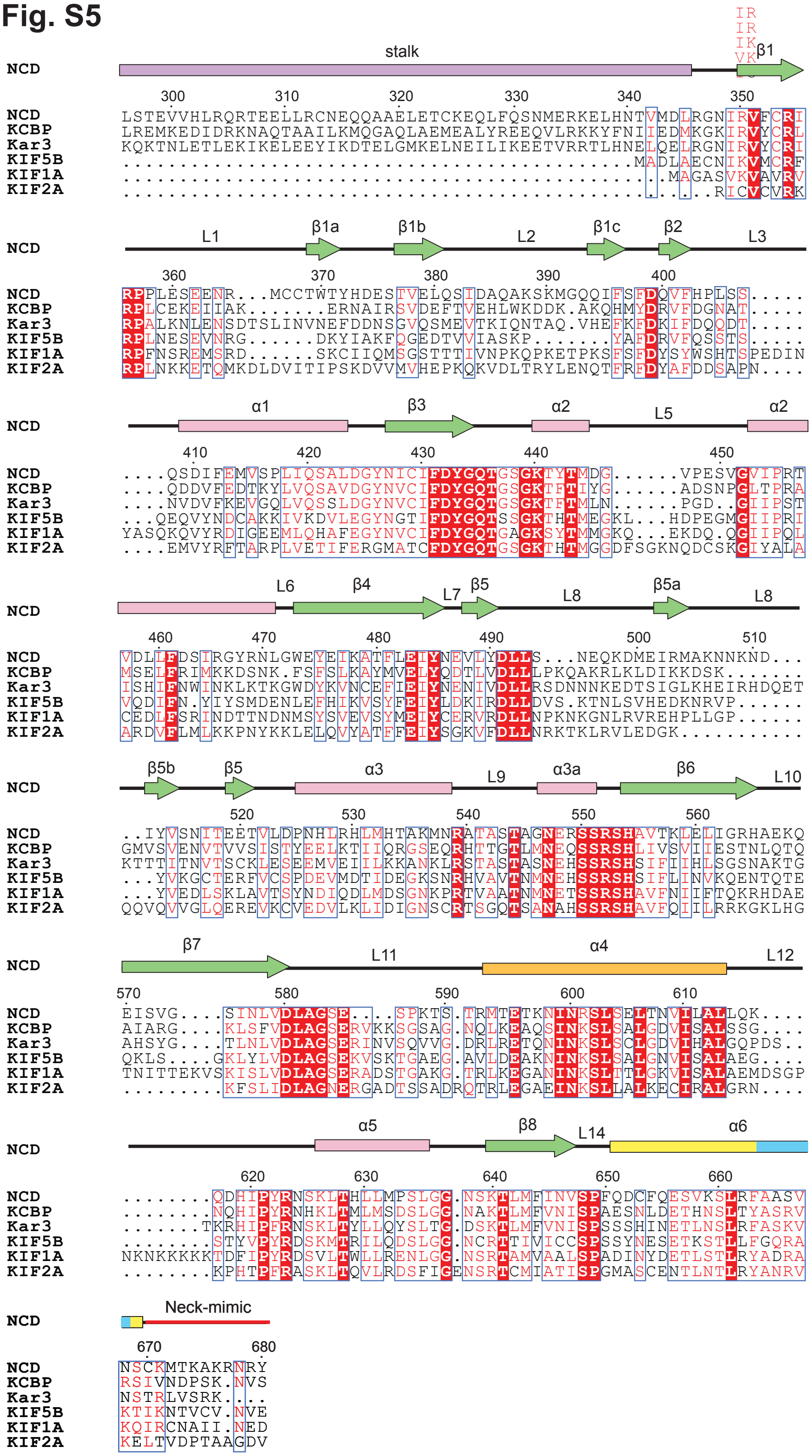


**Fig. S5. Sequence alignment of kinesin-14 and other kinesin family proteins.** Kinesin-14 NCD, KCBP and Kar3 sequence comparison with kinesin-1 KIF5B, kinesin-3 KIF1A and kinesin-13 KIF2A. Kinesin-14 NCD stalk, purple bar; neck-mimic, red line; AASVN motif, cyan bar and shading. Sequence alignment by ENDscript 2.0 (*71*). MT-Ncd NF bound head secondary structure analyzed by DSSP (*72*).

**Fig. S6**

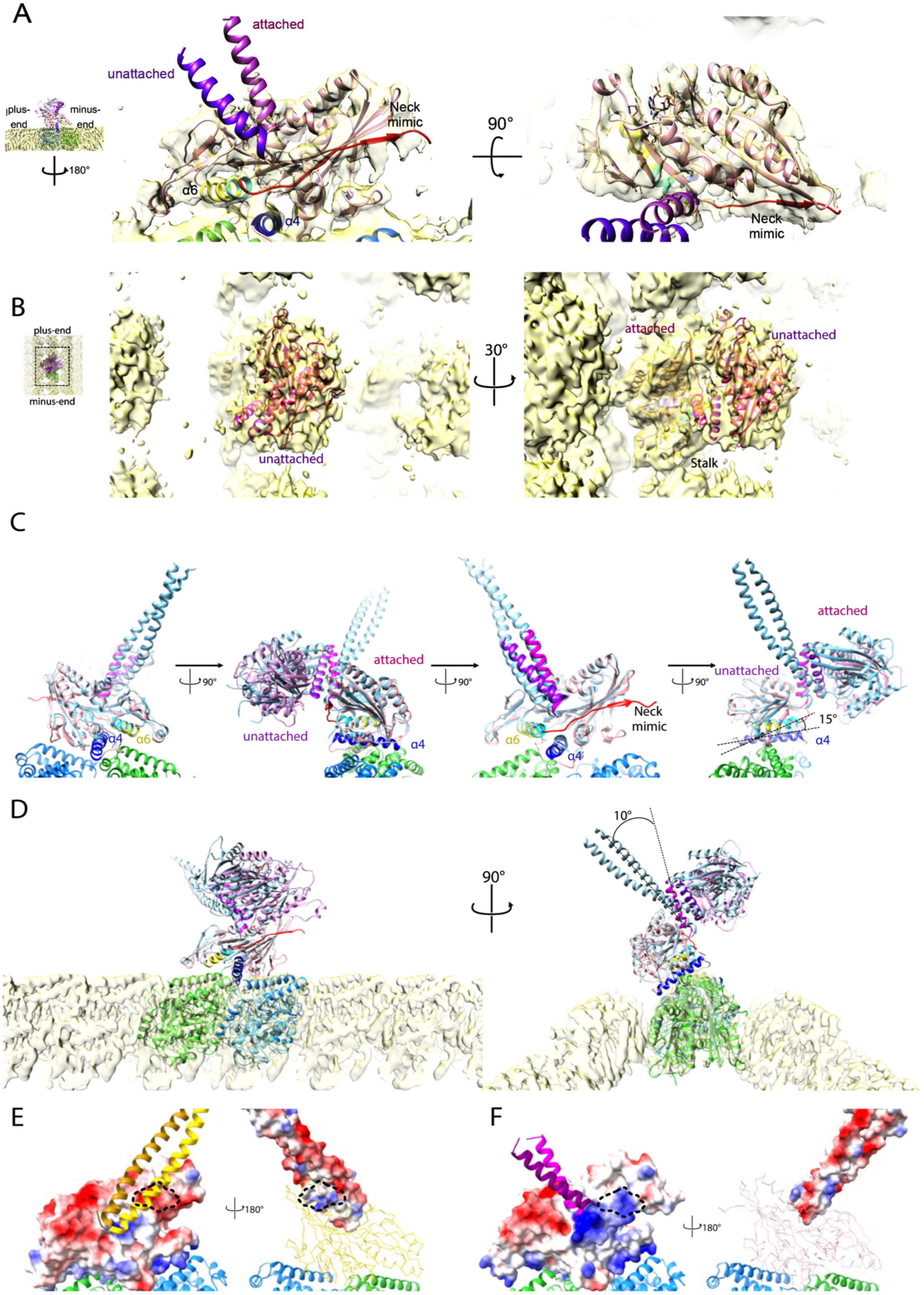
A
attached
unattached
plus-
end
minus-
end
180°
Neck
mimic
α6
α4
90°
Neck
mimic
B
plus-end
minus-end
unattached
30°
attached
unattached
Stalk
C
90°
attached
unattached
α4
α6
90°
α6
α4
Neck
mimic
90°
attached
unattached
15°
α4
D
10°
90°
E
180°
F
180°

**Fig. S6. Structural comparison of MT-Ncd bound and unbound post-PS state.** (**A-D**) MT-Ncd ADP·Pi (ADP-$AlF_3$) cryo-EM structure. The MT-bound motor domain is pale pink and the unattached motor domain is dark pink. Switch II helix α4 is blue, helix α6 is yellow, the helix α6 AASVN motif is cyan, and the neck mimic is red. The stalk of the cryo-EM structure was extended by superimposing the Ncd PDB 1N6M stalk; the stalk helix of the MT-attached head is colored magenta and the stalk helix of the unattached head is purple. (**A**) Unfiltered cryo-EM density map of MT-Ncd ADP·Pi. Inset (left), lower magnification image showing a rotated view from the side with the stalk tilted towards the microtubule minus end. α-Tubulin is green and β-tubulin is blue. (**B**) A 6 Å low pass Gaussian filter was applied to the cryo-EM density. Inset, lower magnification image showing that the view is from the top down and that the microtubule with the bound Ncd motor is oriented vertically with respect to the viewer. (**C**) MT-Ncd ADP·Pi cryo-EM structure superimposed with Ncd 1N6M (pale blue) by aligning MT-attached head of MT-Ncd ADP·Pi and Ncd 1N6M chain B. Viewed from the side, before and after three 90° rotations. (**D**) MT-Ncd ADP·Pi cryo-EM and Ncd 1N6M superposed structure viewed from the side and microtubule minus end. (**E-F**) Electrostatic potential of the MT-attached Ncd head in the (**E**) NF state and (**F**) ADP·Pi state. Red, negative electrostatic potential; blue, positive potential. The black dashed circles indicate (**E**) neck-mimic binding surface (i.e., region where neck mimic will dock) or (**F**) neck-mimic surface after docking.

**Fig. S7**

A

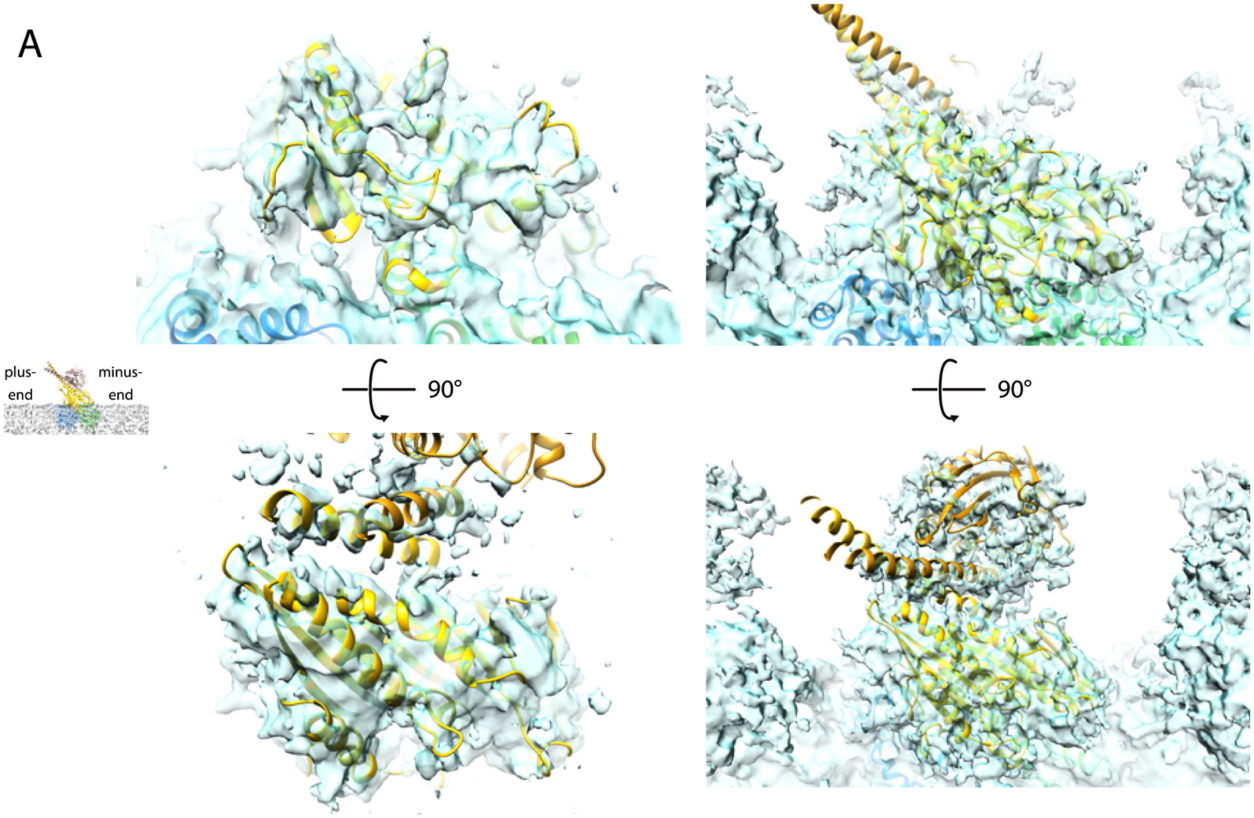


B

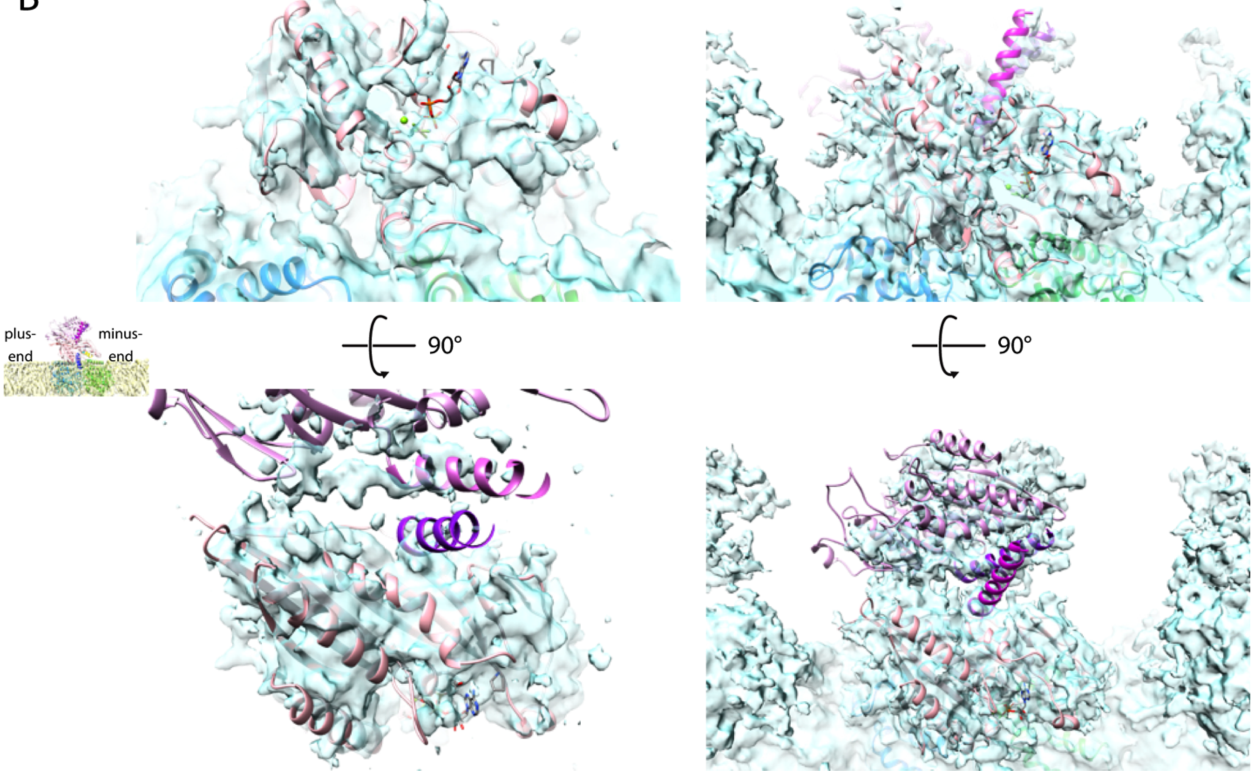

**Fig. S7. Cryo-EM structure of MT-Ncd in the ATP state.** (**A,B**) Cryo-EM density of Ncd in the ATP (AMP·PNP-bound) state fit with (**A**) the MT-Ncd NF (nucleotide free) cryo-EM model (yellow) or (**B**) the MT-Ncd ADP·Pi (ADP-Al$F_3$ bound) cryo-EM model (pink with magenta stalk) viewed from the side (top) or top down (bottom). Insets, lower magnification side view images showing the tilt of the stalk in the (**A**) MT-Ncd NF or (**B**) MT-Ncd ADP·Pi cryo-EM model. The threshold level is set high (0.0007) in panels with a focus on the MT-attached head (left) and low (0.0005) in panels with a focus on the Ncd dimer (right).

### Fig. S8

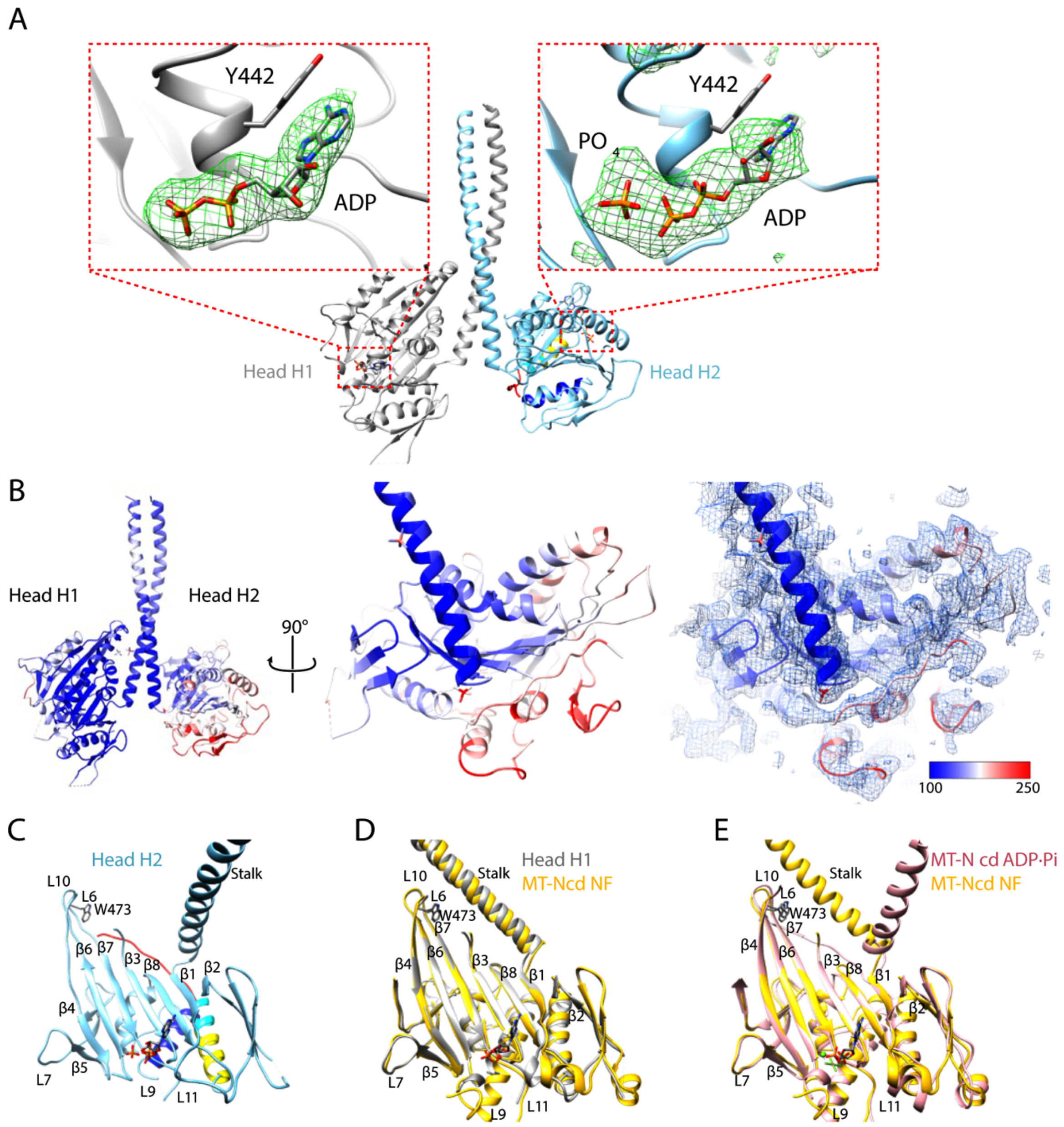


**Fig. S8. MT-unbound Ncd post-PS crystal structure.** (**A**) NcdY485K crystal structure. Insets, head H1 (left) and H2 (right) active sites showing Fo-Fc difference maps (green mesh) of the bound nucleotide with ADP or ADP + free Pi (stick models), respectively, built into the density. Maps contoured at 2.5 σ. The adenine ring of the bound ADP shows ring-stacking interactions with Y442 of the P loop at the nucleotide-binding cleft in both heads. (**B**) NcdY485K crystal structure colored by B-factor (left, center). Scale bar, B-factor range =100-250 $Å^2$. Right, structural elements of high B factor are enclosed by the electron density (2Fo-Fc difference map, blue mesh). (**C**) Central β-sheet structure of head H2 with the stalk helix. Strand topology of the central β-sheet from the outer edge of the head is β54673812. (**D**) Superposition of unbound (white, Ncd crystal structure head H1) and MT-bound (yellow, MT-Ncd NF) pre-PS heads. (**E**) Superposition of MT-bound pre-PS (yellow, MT-Ncd NF) and post-PS (pink, MT-Ncd ADP·Pi) heads.

### Fig. S9

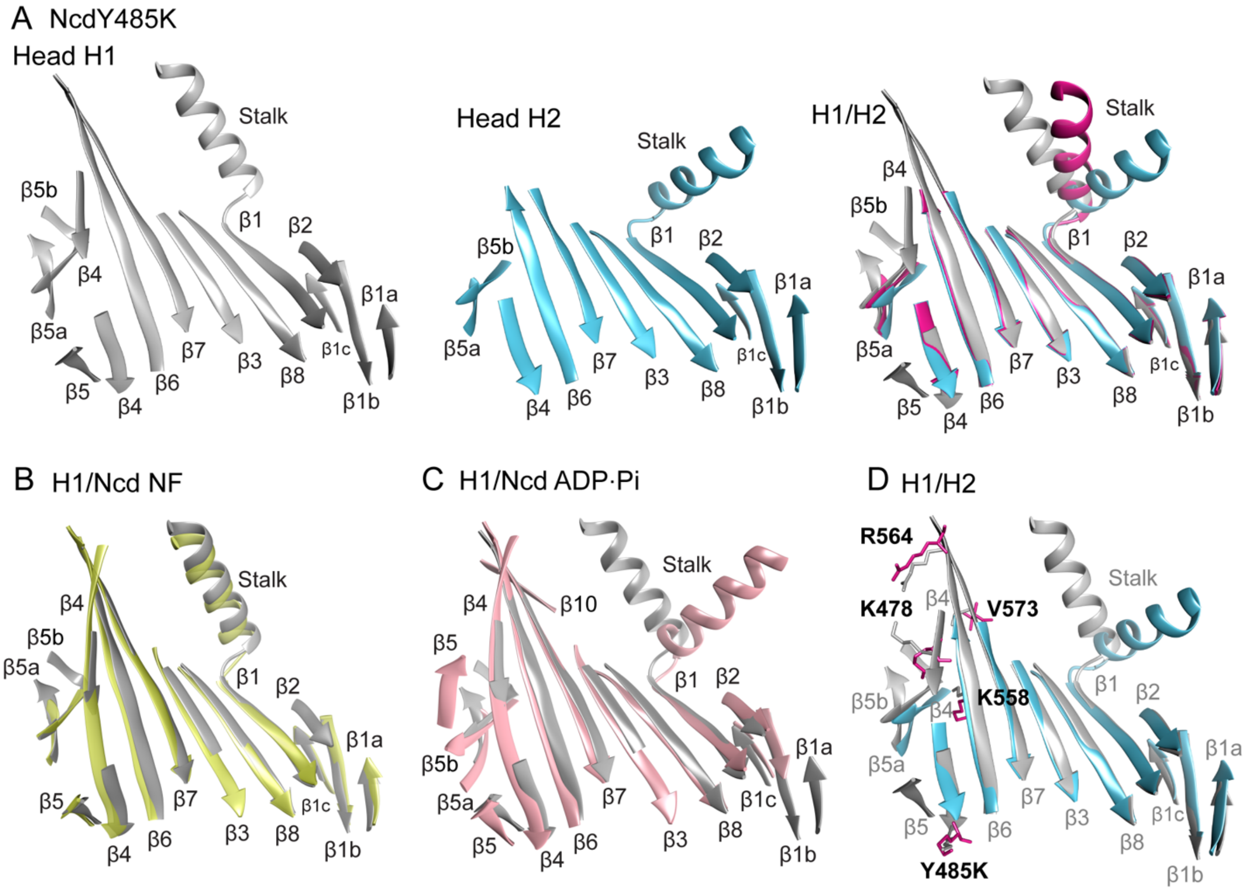


E NcdY485K head H1 vs H2 central β-sheet all atom residue RMSD

RMSD 3.5 Å – 0.2 Å

| | β5 | β4 | β6 | β7 | β3 | β8 | β1 | β2 | β1c | β1b | β1a |
|---|---|---|---|---|---|---|---|---|---|---|---|
| | | | | E570 1.973 | | | | | | | |
| | | | R564 2.679 | | | | | | | | |
| | | E476 1.462 | G563 1.114 | S572 1.175 | | | | | | | |
| | | I 477 1.138 | | V573 2.262 | | | | | | | |
| | T521 0.255 | K478 3.387 | L561 1.596 | G574 1.178 | I 428 0.670 | K640 0.544 | | | | | |
| | E520 1.408 | A479 0.707 | E560 0.969 | S575 1.199 | C429 1.119 | T641 1.456 | I 349 0.975 | | | | |
| | | | L559 0.475 | I 576 0.817 | I 430 0.281 | L642 1.567 | R350 1.625 | | | | |
| | | F481 0.799 | K558 2.307 | N577 0.901 | F431 0.906 | M6430.977 | V351 1.349 | Q400 1.493 | | | |
| | D491 0.833 | L482 0.558 | T557 0.865 | L578 0.964 | A432 0.661 | F644 1.211 | F352 1.255 | V401 0.655 | | | |
| | | E483 0.984 | V556 0.883 | V579 0.707 | Y433 0.803 | I 645 0.913 | C353 0.679 | F402 1.514 | S397 1.067 | T377 0.852 | |
| | | I 484 0.505 | A555 0.525 | D580 0.862 | | N646 1.153 | R354 0.605 | | F396 0.934 | V378 0.906 | Y372 1.897 |
| | V488 1.709 | K485 2.357 | H554 0.314 | | | V647 0.367 | I 355 1.698 | | I 395 1.196 | E378 0.955 | T371 1.356 |
| | | | | | | | | | | L380 1.324 | W370 0.960 |
| | | | | | | | | | | Q381 1.673 | T369 0.991 |
| Strand RMSD$^2$ (Å$^2$) | 44.302 | 205.658 | 175.500 | 122.029 | 30.654 | 79.136 | 96.565 | 48.264 | 27.878 | 58.274 | 75.845 |

Total β-Sheet RMSD$^2$ 964.106 Å$^2$

**Fig. S9. Ncd central β-sheet twisting or distortion.** (**A**) NcdY485K crystal structure central β-sheet, including strands β1a,b,c and β5a,b. Left, NcdY485K head H1 ADP (gray, pre-PS). Middle, NcdY485K head H2 ADP + free Pi (cyan, post-PS). Right, images from morph between H1 and H2 (Chimera *Morph Conformation*; see movie S3) showing β-strand twisting and length changes between the two nucleotide states (dark red, intermediate). (**B**) Ncd central β-sheet movements following microtubule binding and (**C**) the power stroke. (**B**) Superposition of unbound (NcdY485K crystal structure H1, gray) and MT-bound (MT-Ncd NF, yellow) pre-PS heads (movie S4). (**C**) Superposition of unbound (NcdY485K crystal structure H1, gray) and MT-bound post-PS (MT-Ncd ADP·Pi, pink) heads (movie S5). (**D**) NcdY485K β-sheet residues (stick models, bold labels) that undergo the largest movements (>2 Å) between NcdY485K head H1 ADP (pre-PS) and head H2 ADP + free Pi (post-PS) (movie S3). (**E**) Heat map showing NcdY485K head H1 vs H2 central β-sheet all-atom residue RMSD values. Left, scale bar (0.2-3.5 Å RMSD). Bottom, total β-strand and β-sheet $RMSD^2$ (sum of squares) values. NcdY485K head H1 vs H2 mean β-strand RMSD = 1.250±0.230 Å (mean±SD, n=11 β-strands). By contrast, wild-type Ncd 1CZ7 dimer 2 (chain C vs D) heads, both in the ADP pre-PS state, showed a mean β-strand RMSD=0.342±0.082 Å (n=11 β-strands), differing significantly from NcdY485K (P<0.0001, unpaired t test).

### Fig. S10

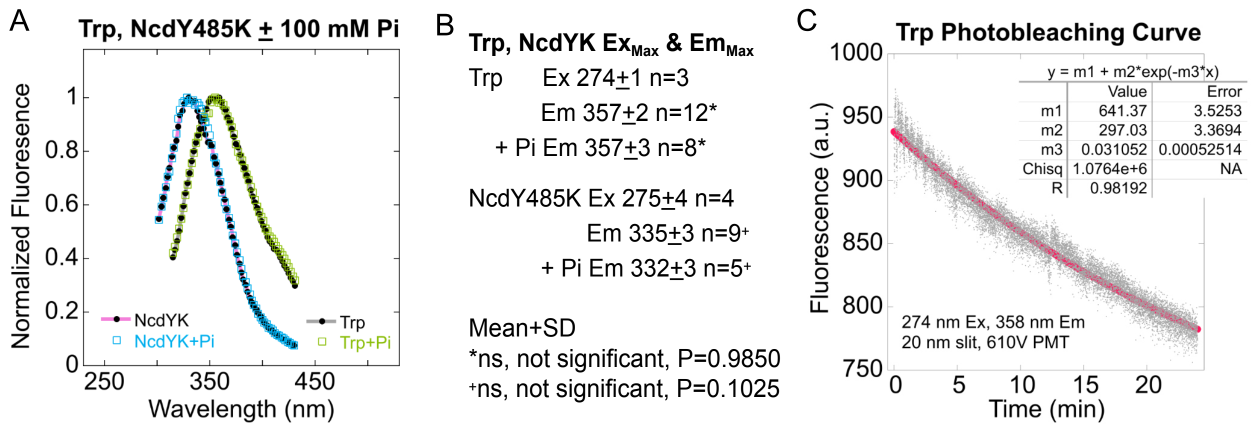


**Fig. S10. Fluorescence assays for NcdY485K structural changes.** (**A**) NcdY485K intrinsic tryptophan (Trp) and free Trp fluorescence emission spectra (Ex, 274 nm) without added NaPi (Left, NcdY485K, pink curve fit; Right, free Trp, gray curve fit) and with 100 mM NaPi (NcdY45K, cyan circles; free Trp, green circles). The spectra without and with 100 mM NaPi overlap for both NcdY485K and free Trp. (**B**) Maximum excitation values for free Trp and NcdY485K are overlapping. Maximum emission values without or with 100 mM NaPi (Pi) do not differ significantly for free Trp (P=0.9820, unpaired t test) or NcdY485K (P=0.1025). However, free Trp $Em_{Max}$ (357±2, n=20) differs significantly from NcdY485K $Em_{Max}$ (333±3, n=14; P<0.0001). Ex, Excitation; Em, Emission; Max, Maximum. (**C**) Free Trp photobleaching curve used to correct for fluorescence loss due to photobleaching in fluorimeter assays. Fits of the data to a single exponential decay model gave a photobleaching decay constant, k=0.00051753/s (correlation coefficient, R=0.98). V, voltage.

**Fig. S11**

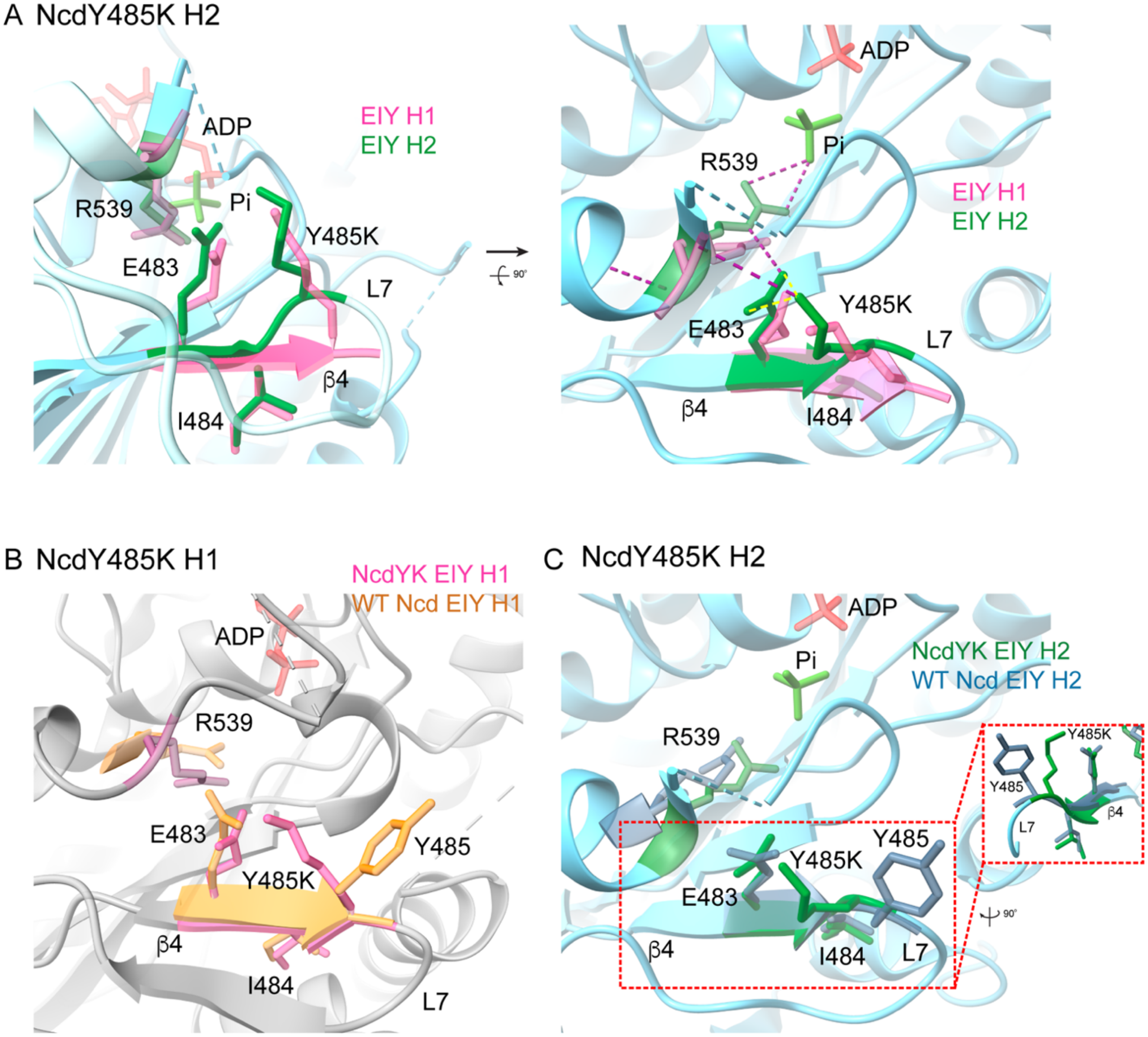


**Fig. S11. NcdY485K mutant effects.** (**A**) NcdY485K head H2 (cyan) showing the H1/H2 superposed EIY motif (H1, pink; H2, green) at the mutation site. Left, Y485K and E483 tilt towards each other in both heads and Y485K has rotated towards E483 in H2. H2 strand β4 is twisted and the distal end has transitioned to a loop. Right, head H2 Y485K is close enough to E483 to form H-bonds (dashed yellow lines) and both residues interact with switch I R539. R539 in head H2 has moved towards the free Pi (yellow green) bound to the H2 active site and is close enough to form H-bonds (dashed purple lines). ADP, pale red. Dashed cyan lines, disordered residues. (**B**) NcdY485K head H1 (gray) with EIY motif (pink) superposed with wild-type (WT) Ncd (PDB 5W3D) head H1 EIY motif (gold). Y485 is tilted away from E483 in WT Ncd head H1. (**C**) NcdY485K head H2 (cyan) with EIY motif (green) superposed with WT Ncd head H2 EIY motif (slate blue). Y485 is tilted away from E483 in WT Ncd head H2. Inset, NcdY485K H2 strand β4 twisting is enhanced compared to WT Ncd β4. The enhanced NcdY485K β-strand twisting is also observed in the increased mean $C_\alpha$-$C_\alpha$ RMSD when the NcdY485K H1 β-sheet is compared to the same NcdY485K H2 (82 atom pairs, 1.154 Å; table S2) or WT Ncd 5W3D H2 (69 atom pairs, 0.614 Å) residues.

**Table S1. Cryo-EM data collection, model building, and refinement statistics**

| | Nucleotide free state | AMP-PNP state | ADP-AlF$_3$ state |
|---|---|---|---|
| **Data collection and processing** | | | |
| **EM equipment** | JEOL CRYO ARM 300 | JEOL CRYO ARM 300 | JEOL CRYO ARM 300 |
| Voltage (kV) | 300 | 300 | 300 |
| Detector | Gatan K3 | Gatan K3 | Gatan K3 |
| Magnification | 60000 | 60000 | 60000 |
| Electron exposure ($e^{-}$/Å$^2$) | 50 | 50 | 50 |
| Defocus range (µm) | -1.7 — -1.4 | -1.7 — -1.4 | -1.7 — -1.4 |
| Pixel size (Å) | 0.752 | 0.752 | 0.752 |
| Initial particle images (no.) | 83412 | 98995 | 75633 |

| **Data processing and refinement** | NF 13PF | NF 14PF | AMPPNP 13PF | AMPPNP 14PF | ADP-AlF$_3$ 13PF | ADP-AlF$_3$ 14PF |
|---|---|---|---|---|---|---|
| EMDB | EMD-39664 | EMD-39665 | EMD-39668 | EMD-39669 | EMD-39666 | EMD-39667 |
| PDBID | 8YY2 | 8YY3 | - | - | 8YY4 | 8YY5 |
| Final particle images (no.) | 20390 | 45321 | 29707 | 30431 | 28547 | 24833 |
| Map resolution of (C1) (Å) | 7.39 | 7.52 | 7.52 | 7.39 | 7.69 | 6.53 |
| Map resolution (symmetrized)(Å) | 3.60 | 3.24 | 3.88 | 3.60 | 4.02 | 3.99 |
| Helical rise (Å) | 9.40 | 8.75 | 9.34 | 8.69 | 9.37 | 8.74 |
| Helical twist (°) | -27.69 | -25.74 | -27.69 | -25.73 | -27.69 | -25.74 |
| FSC threshold | 0.143 | 0.143 | 0.143 | 0.143 | 0.143 | 0.143 |
| Initial model used (PDB code) | NF 14PF | 1CZ7 | | | ADP-AlF$_3$ 14PF | 1N6M |
| Model composition | | | | | | |
| Non-hydrogen atoms | 12673 | 12673 | | | 12306 | 12306 |
| Protein residues | 1596 | 1596 | | | 1545 | 1545 |
| Ligands | 3 | 3 | | | 6 | 6 |
| R.m.s. deviations | | | | | | |
| Bond lengths (Å) | 0.004 | 0.003 | | | 0.003 | 0.015 |
| Bond angles (°) | 0.715 | 0.644 | | | 0.712 | 0.793 |
| Validation | | | | | | |
| MolProbity score | 2.02 | 1.96 | | | 2.12 | 2.14 |
| Clash score | 17.42 | 15.86 | | | 19.10 | 19.88 |
| Rotamer outliers (%) | 0 | 0 | | | 0 | 0 |
| Ramachandran plot | | | | | | |
| Favored (%) | 95.96 | 96.20 | | | 95.09 | 95.09 |
| Allowed (%) | 4.04 | 3.80 | | | 4.91 | 4.91 |
| Disallowed (%) | 0 | 0 | | | 0 | 0 |

**Table S2. MT-Ncd compared to Ncd ADP β-sheet $C_\alpha$-$C_\alpha$ RMSD analysis**

| Ncd structure | MT-bound or unbound head | Nucleotide state | Pruned atom pairs (#) | RMSD (Å) | All atom pairs (#) | RMSD (Å) |
|---|---|---|---|---|---|---|
| ADP* | Unbound | Pre-PS | 82 | 0.000 | 82 | 0.000 |
| MT-Ncd ADP·Pi† | Unbound | Pre-PS | 82 | 0.498 | 82 | 0.498 |
| MT-Ncd NF‡ | Bound | Pre-PS | 82 | 0.649 | 82 | 0.649 |
| MT-Ncd NF‡ | Unbound | Pre-PS | 80 | 0.620 | 82 | 0.701 |
| MT-Ncd ADP·Pi† | Bound | Post-PS | 80 | 0.886 | 82 | 0.953 |
| ADP + free Pi§ | Unbound | Post-PS | 72 | 0.713 | 82 | 1.154 |

*NcdY485K 8YUE crystal structure (this report) chain A head H1 β-sheet residues (349-355,369-371,377-381,395-397,400-402,428-431,473–474,477-479,482-485,488-491,502-505,508-514,520-521,554-565,570-580,640-647; 675 atoms, 672 bonds, 82 residues) were compared with the same residues in the structures shown in the table. RMSD mean values were displayed from pairwise $C_\alpha$-$C_\alpha$ comparisons using Matchmaker in ChimeraX v1.6 (*57*).
†MT-Ncd ADP·Pi 14 PF 8YY5
‡MT-Ncd NF 14 PF 8YY3
§NcdY485K 8YUE chain B head H2

**Table S3. X-ray crystallography statistics for data collection and refinement**

| | NcdY485K (PDB 8YUE) |
|---|---|
| **Data collection** | |
| Space group | $C2_1$ |
| Cell Dimensions a, b, c (Å) | 164.51, 67.62, 94.20 |
| α, β, γ (°) | 90, 96.99, 90 |
| Resolution (Å) | 93.5-3.1 (3.32-3.1) |
| Observed reflections | 119171(15508) |
| Unique reflections | 18107 (2747) |
| $R_{pim}$ | 0.138 (1.04) |
| I/σI | 6.9 (0.8) |
| Completeness (%) | 96.1 (82.3) |
| Redundancy | 6.6 (5.6) |
| **Refinement** | |
| Resolution (Å) | 46.75-3.15 (3.35-3.15) |
| No. reflections | 17466 (1460) |
| $^{*}R_{work}/^{\dagger}R_{free}$ | 0.25 (0.44)/0.29 (0.48) |
| No. atoms Proteins | 5811 |
| Ligand/Ion | 74 |
| B-factors (Å$^2$) Proteins | 138.29 |
| Ligand/Ion | 132.36 |
| RMS deviations Bond lengths (Å) | 0.018 |
| Bond angles (°) | 1.59 |
| Ramachandran statistics Favored regions | 92.0 |
| Allowed regions | 8.0 |
| Disallowed regions | 0 |

Values for the highest-resolution shell are given in parentheses.

$^{*}R_{work} = \Sigma| F_{obs} - F_{calc}|/\Sigma|F_{obs}|$ where $F_{calc}$ and $F_{obs}$ are the calculated and observed structure factor amplitudes, respectively.

$^{\dagger}R_{free}$ = as for $R_{work}$, but for 5.0% of the total reflections chosen at random and omitted from refinement.

**Table S4. Ncd β-strand length changes in crystal structures**

| Crystal structure | Chain | ‡NT state | β-strand residues* (#)† β4 | β6 | β7 |
|---|---|---|---|---|---|
| §WT Ncd 1CZ7 | C | Pre-PS | W473-I484 (12) | H554-H565 (12) | Q569-D580 (12) |
| | D | Pre-PS | E474-I484 (11) | H554-H565 (12) | Q569-D580 (12) |
| WT Ncd 5W3D | A | Pre-PS | W473-I484 (12) | H554-H565 (12) | E570-D580 (11) |
| | B | Post-PS | E474-A479 L482-I484 (9) | H554-V556 L561-R564 (7) | **S575-D580 (6)** |
| NcdY485K¶ | A | Pre-PS | W473-E474 I477-A479 L482-K485 (9) | H554-H565 (12) | E570-D580 (11) |
| | B | Post-PS | E476-I484 (9) | H554-I561 (9) | G574-D580 (7) |
| NcdG347D 3U06 | A | Pre-PS | W473-I484 (12) | H554-H565 (12) | E570-D580 (11) |
| | B | Post-PS | E474-Y485 (12) | H554-R564 (11) | I571-D580 (10) |
| NcdN600K 1N6M | A | Pre-PS | W473-I484 (12) | H554-H565 (12) | E570-D580 (11) |
| | B | Post-PS | **E474-A479 (6)** | A555-E560 I562-R564 (9) | **S572-V573 N577-D580 (6)** |
| NcdT436S 3L1C | A | Pre-PS | W473-I484 (12) | H554-H565 (12) | E570-D580 (11) |
| | B | Post-PS | Y475-F481 (7) | T557-G563 (7) | **V573-S575 (3)** |

*β-strand residues of wild-type Ncd 1CZ7 chain C: strand β4, WEYEIKATFLE1 (12 residues); β6, HAVTKLELIGRH (12 residues); β7, QEISVGSINLVD (12 residues)

†#, number of residues in the β-strand; bold font, ≥50% shorter than wild-type Ncd

‡NT, Nucleotide State; Ncd 1CZ7 chain C and D are pre-power stroke (Pre-PS) nucleotide states; the other Ncd structures show chain A (head H1) in a Pre-PS state and chain B (head H2) in a post-power stroke (Post-PS) state.

§WT, wild type

¶NcdY485K 8YUE, this report

**Table S5. Kinesin-14 NcdY485K fluctuations during ATP binding and force production**

| Motor protein | MT/actin binding | MT/actin unbinding | ATP hydrolysis | Power stroke | Recovery stroke | References |
|---|---|---|---|---|---|---|
| Kinesin | -ADP | ADP+Pi | MT-bound | ATP binding<br>Multistate | Pi release<br>Unbound | *12*, this report |
| Myosin | -ADP<br>(ADP+Pi)* | +ATP | Unbound | Pi release<br>Multistate | ADP release<br>Actin-bound | *19* |
| Dynein | -ADP<br>(+ADP)* | +ATP | Unbound | Pi release<br>Multistate | ADP release<br>MT-bound | *69* |

*Rebinding following ATP hydrolysis

### Computer Script

#### Chimera RMSD script for all-atom β-sheet residue differences

A script like the one below was used to obtain the NcdY485K head H1 vs H2 RMSD values shown in the heat map in fig. S9E. Residue numbers for this analysis depend on the criteria used to define the β-strands and were changed to obtain the RSMD values for Ncd 1CZ7 chain C vs chain D β-strand residue differences (see **Materials and methods**).

```
from chimera import runCommand
from chimera.tkgui import saveReplyLog
import numpy as np
#%% Define central beta sheet residues
beta1 = np.linspace(349, 355, 7).astype(int)
beta1a = np.linspace(369,372,4).astype(int)
beta1b = np.linspace(377,381,5).astype(int)
beta1c = np.linspace(395,397,3).astype(int)
beta2 = np.linspace(400, 402, 3).astype(int)
beta3 = np.linspace(428, 433, 6).astype(int)
beta4_1 = np.linspace(476, 479, 4).astype(int)
beta4_2 = np.linspace(481, 485, 5).astype(int)
beta5_1 = [488, 491]
beta5_2 = [520, 521]
beta6_1 = np.linspace(554, 561, 8).astype(int)
beta6_2 = [563,564]
beta7 = [570,572,573,574,575,576,577,578,579,580]
beta8 = np.linspace(640, 647, 8).astype(int)
#Array of all residue numbers
strandArr = [beta1, beta1a, beta1b, beta1c, beta2, beta3, beta4_1, beta4_2, beta5_1, beta5_2,
beta6_1, beta6_2, beta7, beta8]
#%% Run Chimera commands and save outputs
runCommand('mm #0 #1')
for n in range(len(strandArr)):
    strand = strandArr[n]
    for x in range(len(strand)):
        residue = int(strand[x])
        runCommand("rmsd #0:" + str(residue) + " #1:" + str(residue))
#Save outputted reply log as text file
saveReplyLog("NcdY485K atPDB Head A vs B, 06-26-24.txt")
```

**Supplemental Movies**

**Movie S1. Ncd NF | ADP·Pi   Power stroke, β-sheet twisting ($C_\alpha$, ribbon)**
**Movie S2. Ncd ADP·Pi | ADP+free Pi   Docked neck mimic, β-strands to loops**
**Movie S3. Ncd ADP+free Pi | ADP   Recovery stroke, β-sheet refolding**
**Movie S4. Ncd ADP | NF   Stalk movement, β-sheet twisting**
**Movie S5. Ncd ADP | ADP·Pi   Power stroke, β-sheet twisting**
**Movie S6. NcdCycle   Ncd mechanochemical cycle model**
